# Temperature-Dependent Fold-Switching Mechanism of the Circadian Clock Protein KaiB

**DOI:** 10.1101/2024.05.21.594594

**Authors:** Ning Zhang, Damini Sood, Spencer C. Guo, Nanhao Chen, Adam Antoszewski, Tegan Marianchuk, Archana Chavan, Supratim Dey, Yunxian Xiao, Lu Hong, Xiangda Peng, Michael Baxa, Carrie Partch, Lee-Ping Wang, Tobin R. Sosnick, Aaron R. Dinner, Andy LiWang

**Author notes:** Current address: Qingdao Institute of Bioenergy and Process, Chinese Academy of Sciences; Shandong Energy institute; Qingdao New Energy Shandong Laboratory, Qingdao 266101, China (Ning Zhang). Current address: Biogen, Cambridge, MA, United States. Current address: Genentech, San Francisco, CA, United States. Current address: AbCellera, Vancouver, Canada (Lu Hong, Aaron R. Dinner). Co-first authors (Ning Zhang, Damini Sood, Spencer C. Guo).

## Abstract

The oscillator of the cyanobacterial circadian clock relies on the ability of the KaiB protein to switch reversibly between a stable ground-state fold (gsKaiB) and an unstable fold-switched fold (fsKaiB). Rare fold-switching events by KaiB provide a critical delay in the negative feedback loop of this post-translational oscillator. In this study, we experimentally and computationally investigate the temperature dependence of fold switching and its mechanism. We demonstrate that the stability of gsKaiB increases with temperature compared to fsKaiB and that the Q10 value for the gsKaiB → fsKaiB transition is nearly three times smaller than that for the reverse transition. Simulations and native-state hydrogen-deuterium exchange NMR experiments suggest that fold switching can involve both subglobally and near-globally unfolded intermediates. The simulations predict that the transition state for fold switching coincides with isomerization of conserved prolines in the most rapidly exchanging region, and we confirm experimentally that proline isomerization is a rate-limiting step for fold switching. We explore the implications of our results for temperature compensation, a hallmark of circadian clocks, through a kinetic model.

## INTRODUCTION

The fold of a protein is the *(i)* overall three-dimensional arrangement of secondary structures (i.e., architecture) and *(ii)* the path of the polypeptide chain through the structure (i.e., topology) (1). An exciting finding over the last several years is that some proteins do not follow the classic “one sequence, one fold” paradigm but can switch between different folds reversibly under physiological conditions (2, 3). These proteins belong to the relatively new class of so-called metamorphic proteins (4).

One of the most well-studied metamorphic proteins is KaiB, which is a core component of the oscillator of the cyanobacterial circadian clock (5). Circadian clocks are an adaptation to daily oscillations in ambient light and temperature and play important roles in the fitness and health in diverse organisms (6, 7). KaiB, together with KaiA, generates a circadian rhythm of phosphorylation in the clock protein, KaiC, to regulate downstream expression of clock-controlled genes (8).

In cyanobacteria with a circadian clock, KaiB exists in an equilibrium between two distinct folds with vastly different stabilities. The stable fold of KaiB with secondary structure βαββααβ is found only in KaiB homologs, and we refer to it as ground-state KaiB, or gsKaiB (9–12). The fold that binds KaiC is otherwise unstable; it is a thioredoxin-like fold with secondary structure βαβαββα, and we refer to it as fold-switched KaiB, or fsKaiB (Error! Reference source not found.). The much higher stability of gsKaiB relative to fsKaiB is the reason why the latter was not discovered until a decade after the structure of gsKaiB was solved by X-ray crystallography.

While fsKaiB is monomeric, gsKaiB self-associates to form dimers which in turn come together to form an asymmetric dimer of dimers. Thus, prior to binding KaiC, homotetramers of gsKaiB likely need to dissociate into monomers and then undergo a gsKaiB → fsKaiB fold switch (13, 14). Due to the instability of fsKaiB, it binds to KaiC slowly, on the hours time scale (15–17). Mutations that enhance the stability of fsKaiB relative to gsKaiB increase the rate of KaiB-KaiC binding but also abrogate clock function, suggesting that the slowness of binding provides an important delay in the negative feedback loop of this clock (5). Each evening when fsKaiB binds KaiC, it inactivates KaiA (13, 15), displaces the sensor histidine kinase SasA from KaiC (15), activates the phosphatase activity of CikA (18), and binds the protein KidA, which tunes the period of the KaiABC oscillator (19). Thus, fsKaiB is a hub for nighttime signaling events.

Despite the importance of KaiB fold switching to the cyanobacterial circadian clock, much remains unknown about the mechanism of fold switching. Here, we describe experimental and computational evidence that the fold-switching thermodynamics and kinetics are sensitive to temperature. We then demonstrate that KaiB can undergo both subglobal and global unfolding during fold switching and that isomerization of three conserved prolines located in a dynamic region of about ten residues is rate-limiting. We discuss the implications of our results for temperature compensation.

## RESULTS

### The gsKaiB ⇌ fsKaiB equilibrium is sensitive to temperature

Our studies utilize KaiB from the thermophile *Thermosynechococcus elongatus (vestitus) BP-1*, but we expect the results to also apply to KaiB from the widely studied mesophile *Synechococcus elongatus* PCC 7942 because the periods of the oscillators from both organisms are relatively insensitive to temperature (20, 21) and key residues are conserved. Earlier, we found that truncating *T. elongatus* KaiB after residue 94 and introducing Y8A, D91R (or G89A), and Y94A amino acyl substitutions shifts the gsKaiB ⇌ fsKaiB equilibrium to the right such that around room temperature the fsKaiB and gsKaiB states are similarly populated (5) (**Figs. S1 and S2**). Henceforth, we refer to *T. elongatus* KaiB 1-94 Y8A, D91R, Y94A as KaiB^D91R^ (and KaiB 1-94 Y8A, G89A, Y94A as KaiB^G89A^).

KaiB^D91R^ (and KaiB^G89A^) were used to characterize the temperature dependence of the gsKaiB ⇌ fsKaiB equilibrium. Size-exclusion chromatography indicated that the KaiB^D91R^ construct is monomeric at room temperature (**Fig. S3**). ^15^N,^1^H heteronuclear single-quantum correlation (HSQC) NMR spectroscopy of ^15^N-enriched KaiB^D91R^ (and KaiB^G89A^) revealed that the gsKaiB ⇌ fsKaiB equilibrium is sensitive to temperature (**Fig. 2a**). ^15^N,^1^H HSQC spectra of KaiB^D91R^ (and KaiB^G89A^) were collected at 20, 25, 30, and 35 °C under equilibrium conditions, and Δ*G* values were estimated from the volume (population) ratios of the assignable peaks at each temperature (**Figs. 2b and S4**; see SI). Fitting the data to Δ*G* = Δ*H* – *T*Δ*S* shows that the gsKaiB ⇌ fsKaiB reaction is enthalpically driven (Δ*H* < 0 and Δ*S* < 0). Using the Protein Interactions Calculator (22), we compared intrachain contacts between gsKaiB (1VGL) and fsKaiB (5JWR), considering only residues in common between the two constructs and found that gsKaiB has approximately 16% more nonpolar contacts (and 8% fewer polar contacts). We thus interpret the temperature dependence to result in part from the hydrophobic interactions weakening at low temperature (23, 24); this may also account for temperature-dependent fold switching of a designed protein (25).

**Figure 1.**
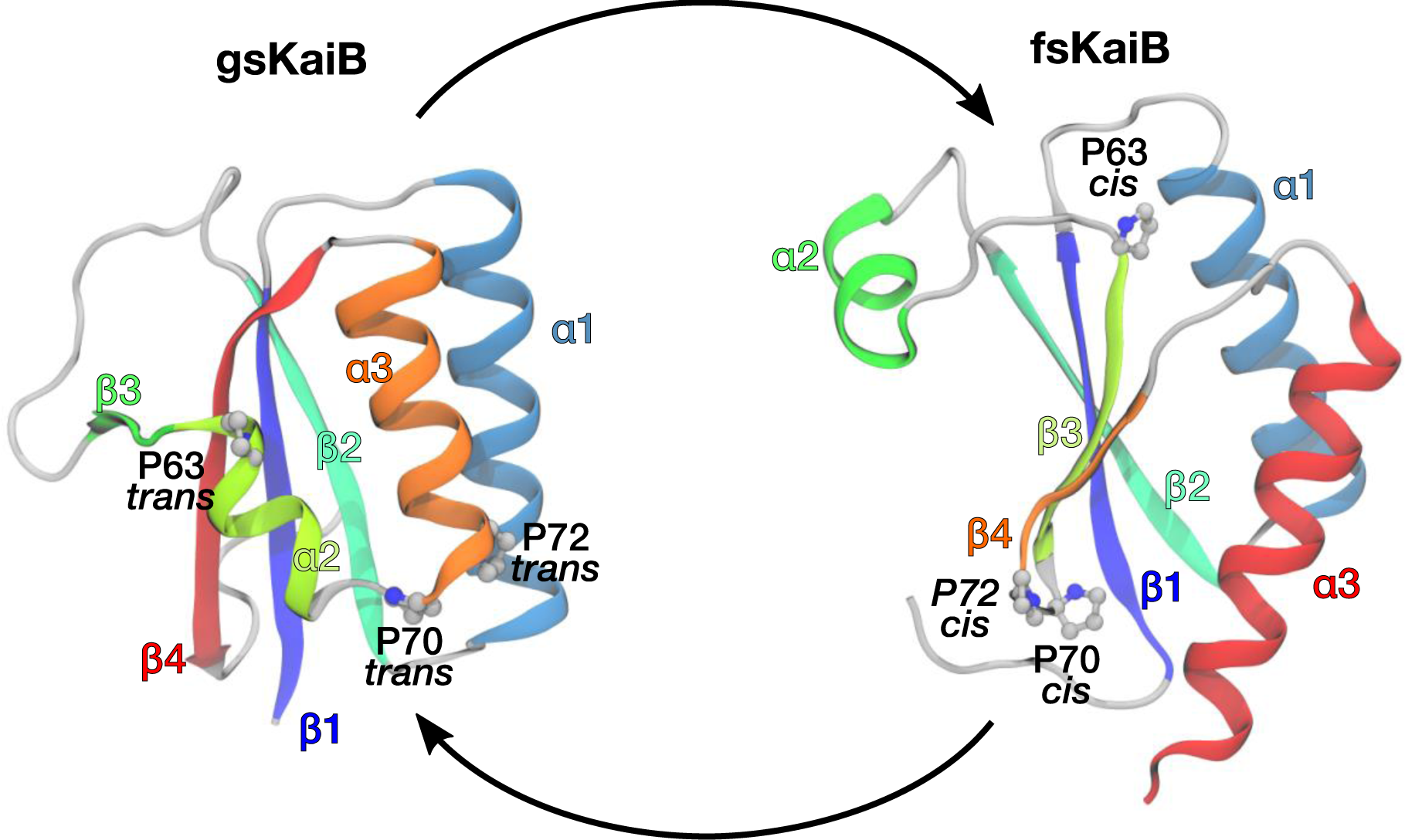
KaiB switches reversibly between two distinctly different folds. Three proline residues — P63, P70, and P72 — are *trans* in the ground-state fold of KaiB (gsKaiB) and *cis* in the fold-switched fold of KaiB (fsKaiB). PDB IDs 1VGL and 5JYT were used in VMD (78) to generate the ribbon diagrams for gsKaiB and fsKaiB, respectively.

**Figure 2.**
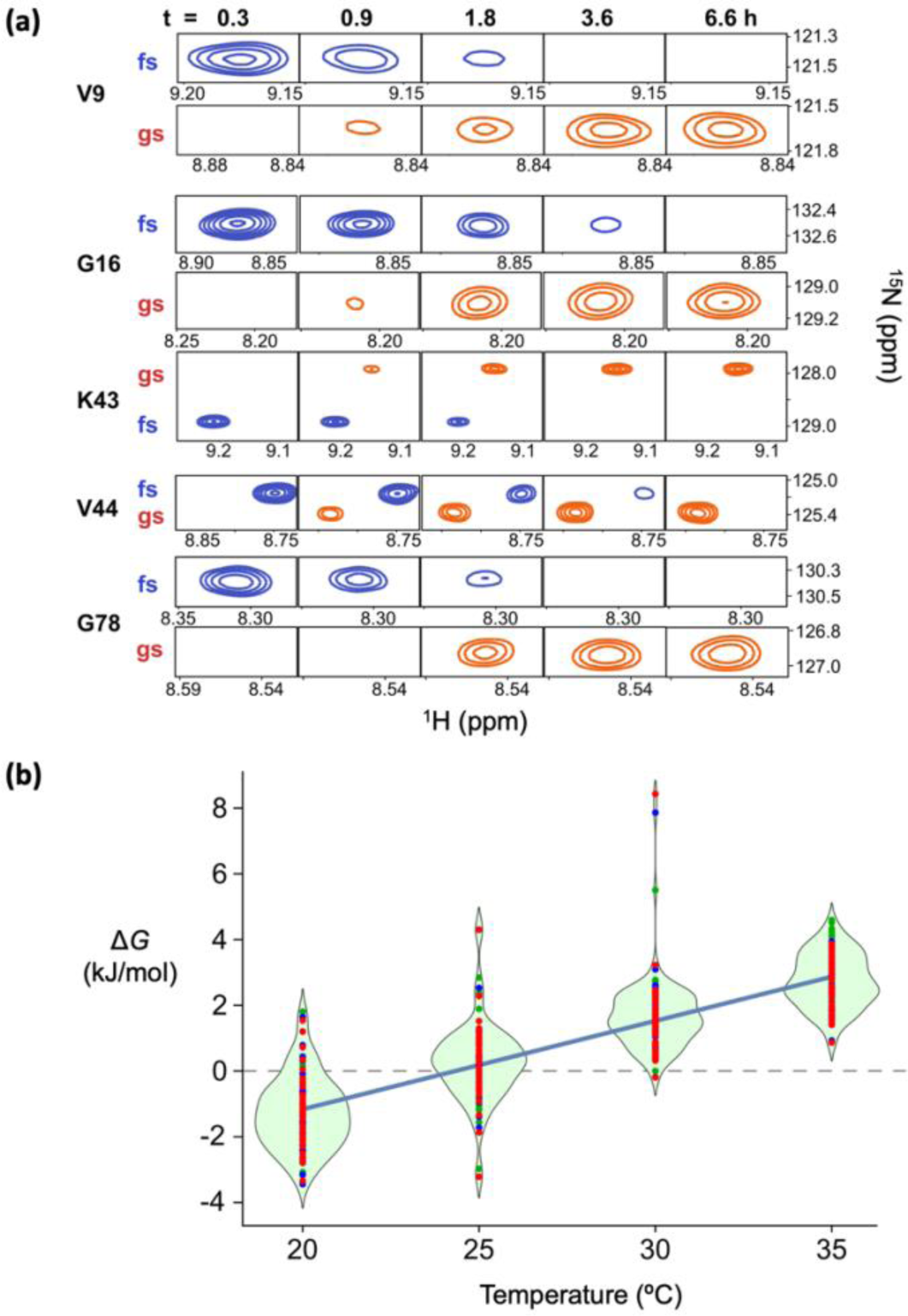
The gsKaiB^D91R^ ⇌ fsKaiB^D91R^ equilibrium is sensitive to temperature. **(a)** A sample of ^15^N-enriched KaiB^D91R^ was incubated at 4 °C for 24 h before inserting it into a 14.1 T NMR spectrometer that was set to 35 °C. ^15^N,^1^H HSQC spectra were collected over time and some representative regions are shown here. **(b)** Residue-specific Δ*G* values for the gsKaiB^D91R^ → fsKaiB^D91R^ reaction are plotted as red, green, and blue points, one color for each replicate, and are superimposed on distribution diagrams for each temperature sampled. The blue line is a fit to mean values using the equation Δ*G* = Δ*H* – *T*Δ*S*. Fitting the data for individual residues yields Δ*H* = −82 ± 18 kJ mol^-1^ and Δ*S* = −276 ± 58 J mol^-1^ K^-1^, indicating that the gsKaiB^D91R^ → fsKaiB^D91R^ reaction is enthalpically driven.

### The activation free energies, Δ*G*^‡^_gs→fs_ and Δ*G*^‡^_gs←fs_, have opposite temperature dependencies

Next, we wanted to determine the extent to which the activation free energies for the forward and reverse fold-switching reactions, Δ*G*^‡^_gs→fs_ and Δ*G*^‡^_gs←fs_, respectively, are dependent on temperature. We incubated ^15^N-enriched KaiB^D91R^ (and KaiB^G89A^) at 4 °C for at least 24 h before collecting HSQC spectra as a function of time at 20 °C, 25 °C, 30 °C, and 35 °C. As shown in **Fig. 3a** for a representative residue, G16, we fit time-dependent changes in fractional populations using integrated volumes of assignable gsKaiB and fsKaiB HSQC peaks of ^15^N-enriched KaiB^D91R^ (and KaiB^G89A^) to obtain the observed rate of fold switching, *k*_obs_, upon a jump in sample temperature. For each temperature jump, the forward and reverse rate constants for fold switching, *k*_gs→fs_ and *k*_gs←fs_, were determined from *k*_obs_ and the equilibrium populations of gsKaiB and fsKaiB for each residue with an assignable set of HSQC peaks. These rate constants were used to calculate Δ*G*^‡^_gs→fs_ and Δ*G*^‡^_gs←fs_ using transition-state theory (see SI for details). As shown in **Figs. 3b** and **S4b,** Δ*G*^‡^_gs→fs_ increases, whereas Δ*G*^‡^_gs←fs_ decreases with temperature. An equivalent perspective can be obtained by calculating Q10 = *k_T_*_+10 °C_/*k_T_*, where *k_T_* is the rate constant at temperature *T* (**Fig. 3c**). Using measurements at 25 °C and 35 °C, Q10 values for *k*_gs→fs_ and *k*_gs←fs_ are 2.1 ± 0.3 and 5.7 ± 0.8, respectively, for KaiB^D91R^. Consequently, the rate of the gsKaiB → fsKaiB transition increases considerably less with temperature than the rate of the reverse reaction does.

**Fig. 3.**
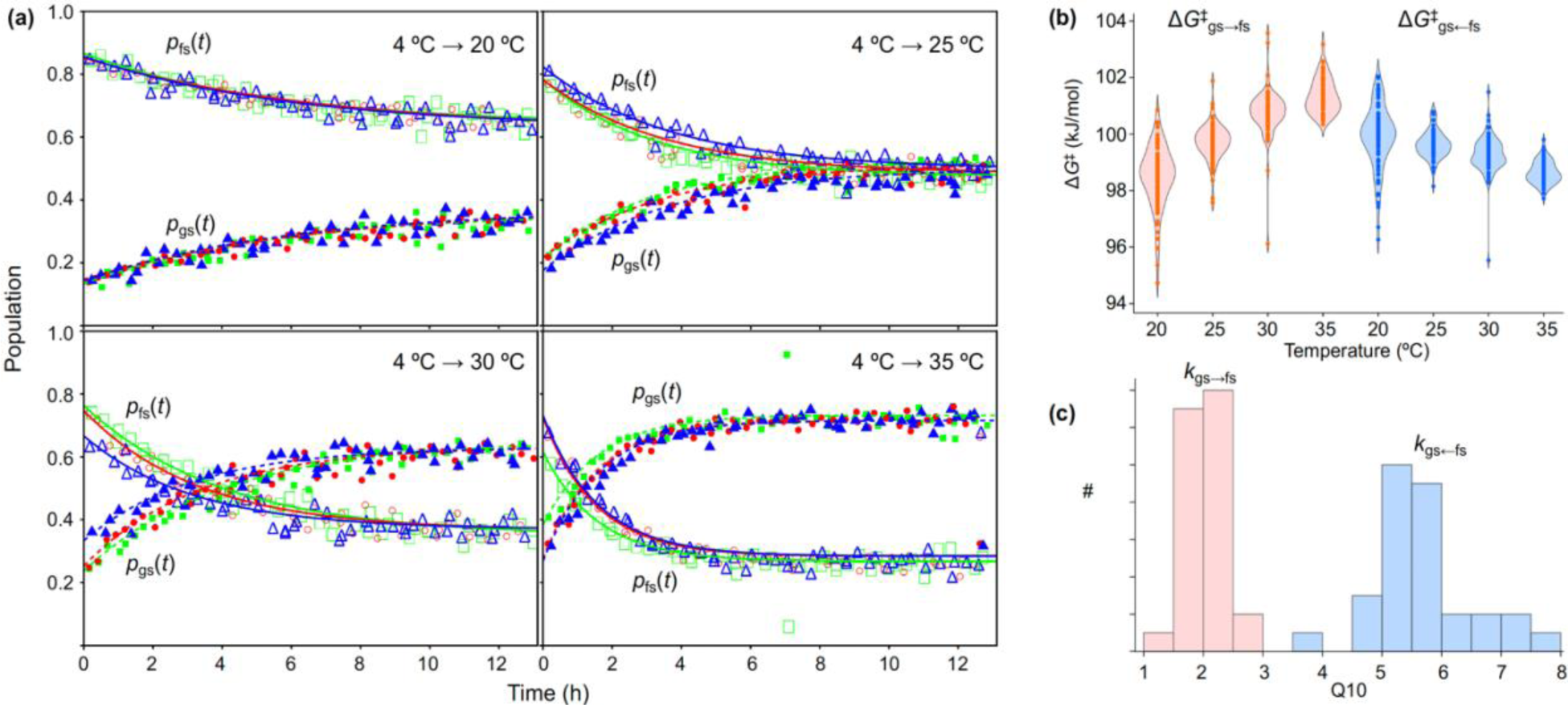
The kinetics of gsKaiB^D91R^ → fsKaiB^D91R^ and gsKaiB^D91R^ ← fsKaiB^D91R^ fold switching differ in their dependence on temperature. **(a)** Kinetics of gsKaiB^D91R^ ← fsKaiB^D91R^ fold switching of residue G16 under different temperature jumps. A sample of ^15^N-enriched KaiB^D91R^ was incubated at 4 °C for at least 24 h before inserting it into a 14.1 T NMR spectrometer that was set at either 20 °C, 25 °C, 30 °C, or 35 °C. ^15^N,^1^H HSQC spectra were collected at regular intervals after a dead time of approximately four minutes. Open and solid symbols represent fractional populations of residue G16 in the gsKaiB^D91R^ and fsKaiB^D91R^ folds, respectively. Red, green, and blue colors represent separate experiments, each of which used a freshly prepared sample. HSQC peak volumes were determined by nmrPipe and nmrDraw. **(b)** Δ*G*^‡^_gs→fs_ and Δ*G*^‡^_gs←fs_ as a function of temperature for all residues, and **(c)** histograms of residue-specific Q10 values for *k*_gs→fs_ and *k*_gs←fs_. The forward and reverse reactions are rendered in pink and blue, respectively. In (b) and (c) the three replicates were pooled.

### Simulations reveal a temperature dependent fold-switching mechanism

To interpret these observations, we investigated how temperature affects fold-switching thermodynamics and kinetics using molecular dynamics simulations. While direct simulation of fold switching remains prohibitively costly, we were able to characterize KaiB’s behavior by combining a machine-learned near-atomic resolution model, Upside (26), with a computational framework for estimating thermodynamics and kinetics from many short simulations that sample portions of the fold-switching transition (27, 28).

We simulated fold switching of KaiB^D91R^ at three temperatures ranging from a temperature where both gsKaiB and fsKaiB structures are stable to near the melting temperature (**Upside temperature calibration** section of Supplementary Information). The computed Δ*G*_gs→fs_ was 3–6 *k*_B_*T* (**Figs. 4a** and **S5**), and gsKaiB is stabilized with respect to fsKaiB at higher temperatures, consistent with the experimental trend. The Upside Δ*G*_gs→fs_ values are somewhat larger than the experimentally observed ones. For comparison, we also used all-atom MD simulations to compute the temperature dependence of the fold-switching free energy difference from the temperature dependence of the free energy of each individual fold (see SI). The relative stabilization from these simulations was approximately 4 kJ/mol (**Fig. S5**), consistent with the experimental findings. Based on this comparison, we infer that the Upside temperature range is somewhat higher and broader than that studied experimentally.

**Figure 4.**
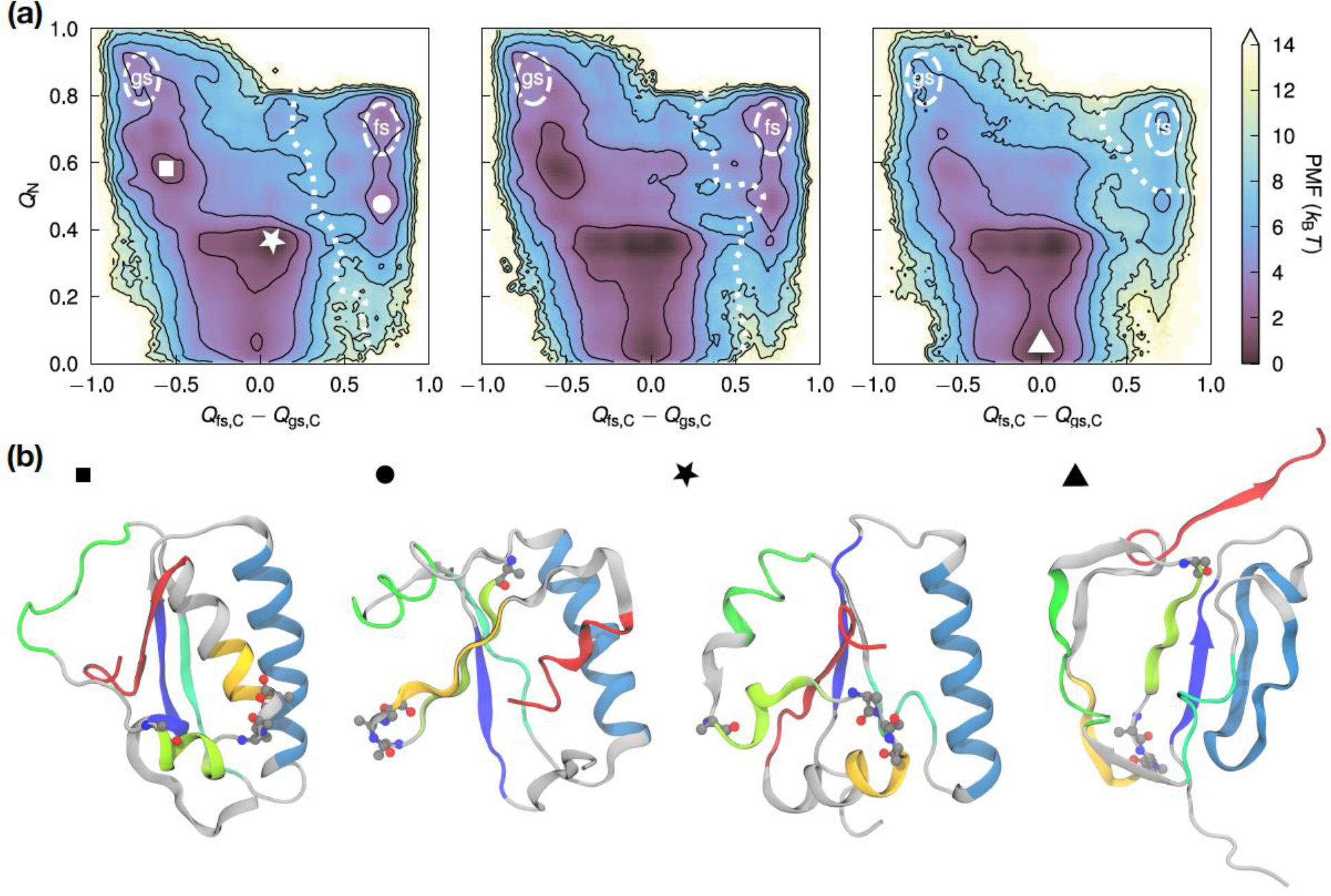
Fold switching in an Upside model of KaiB^D91R^ involves subglobally and globally unfolded intermediates. **(a)** Potential of mean force as a function of the fraction of N-terminal contacts (*Q*_N_) and the difference in C-terminal contacts to the fs and gs structures (*Q*_fs,C_ – *Q*_gs,C_). From left the right, the Upside temperature is increasing. Contour lines are drawn every 2 *k*_B_*T*. The approximate location of the transition state (*q* = 0.5) is marked as a dashed line. **(b)** Selected structures corresponding to CV values in the PMF in **(a)**. The sequence is colored by the position of secondary structures in fsKaiB and P63, P70, P71, and P72 are shown as balls and sticks.

Our analysis for the Upside simulations enables us to compute an important statistic for dissecting the fold switching mechanism: the probability of reaching the fsKaiB state before the gsKaiB state from a given conformation (*q*). This statistic serves as a measure of progress (i.e., reaction coordinate), and we define the transition state as *q* = 0.5. We interpret *q* structurally using the fraction of native contacts (29) in the N-terminal half of the protein (*Q*_N_) and the difference in the fractions of C-terminal contacts between the gsKaiB and fsKaiB states (*Q*_fs,C_ – *Q*_gs,C_; **Table S1**). The transition state cuts diagonally at *Q*_fs,C_ – *Q*_gs,C_ ≈ 0.5 (**Fig. S6**) and shifts toward the fsKaiB state as the temperature increases, consistent with the Hammond postulate (30–32).

Potentials of mean force (free energies) as functions of these coordinates are shown in **Figs. 4a** and **S7**. There is a barrier between the unfolded intermediates and the fsKaiB state, consistent with the location of the transition state, as well as metastable intermediates at (*Q*_fs,C_ – *Q*_gs,C_, *Q*_N_) ≈ (–0.6, 0.6), (0.0, 0.4), (0.0, 0.0), and (0.7, 0.5), for which we show selected structures **Fig. 4b**). Structures with the C-terminal half folded and the N-terminal half unfolded are too high in free energy to be accessed, as indicated by the absence of points in the lower left and right corners of the plots in **Fig. 4a**.

The intermediates marked with a square, star, and circle are on the fold-switching pathway, corresponding to loss or gain of individual C-terminal secondary structures. We show potentials of mean force as functions of CVs characterizing individual secondary structure transitions (see **Collective variables** section of Supplementary Information) in **Fig. S8**. The presence of multiple minima in these plots indicates that these secondary structures can generally transition independently. At the same time, the diagonal symmetry of the *Q*_fs,β4_ *– Q* _gs,α3_ and *Q*_fs,α3_ *– Q* _gs,β4_ plot suggests that the α2_gs_ ⇄ β3_fs_ and α3_gs_ ⇄ β4_fs_ regions form the β3_fs_/β4_fs_ sheet together, while the lack of an intermediate at (1, –1) in the *Q*_fs,β4_ *– Q* _gs,α3_ and *Q*_fs,β3_ *– Q*_gs,α2_ plot suggests that switching β4_gs_ to α3_fs_ is required before switching to β4_fs_ (and β3_fs_). The intermediate marked with a triangle in **Fig. 4** is off-pathway and corresponds to full unfolding of the N-terminal half of the protein. As one would expect, increasing the temperature stabilizes this globally unfolded intermediate.

The picture that emerges is that fold switching can proceed by pathways in which the N-terminal half of the protein unfolds to varying degrees, with higher temperatures favoring a globally unfolded intermediate. Because this intermediate is on the gs side of the transition state (*q* ≤ 0.3, **Fig. S6**), the reactive flux for fold switching involving this intermediate (**Reactive flux through unfolded state** section of Supplementary Information) decreases from 75% to 60% in the gs→fs direction as temperature increases, while the opposite is true for the gs←fs direction (i.e., in both cases, the system tends toward the intermediate and, in turn, gsKaiB as temperature increases, **Fig. S7**). Given that the Upside temperature range is likely higher than the experimental one, as noted above, the global unfolding pathway may be overemphasized by the Upside simulations. Nevertheless, they suggest that KaiB can transition by either global or subglobal unfolding.

### Hydrogen-deuterium exchange experiments and simulations are consistent with large-scale unfolding for a fs variant of KaiB

To test the predictions of the simulations that KaiB can transition by either global or subglobal unfolding, we probed fsKaiB’s free energy surface with native-state hydrogen-deuterium exchange (HDX) experiments. HDX experiments performed on proteins under equilibrium conditions can provide insights into regions of proteins that undergo thermally induced local, subglobal, and global unfolding events by quantifying the sensitivity of HDX rates to the addition of low-to-moderate amounts of denaturant (33). HDX rates of residues whose exchange is determined by local unfolding events can be identified by their having minimal denaturant dependence while the rates of residues that undergo HDX upon larger-scale subglobal or global unfolding events have higher sensitivity (large *m*-values).

Unfortunately, not all peaks in the HSQC spectrum for the α2-β3-β4 region in KaiB^D91R^ could be assigned. Therefore, we created a construct for HDX experiments that is more stable in the fsKaiB fold — *T. elongatus* KaiB 1-99 Y8A P71A G89A D91R Y94A — hereafter fsKaiB^HDX^. For this variant, resonances in the α2-β3-β4 region were assignable. We measured HDX rates at 50+ sites at pH 5.5 and 30+ sites at pH 6.5 at urea concentrations of 0, 0.5, 1.0, 1.5, 2.0, and 2.5 M urea. The most stable regions were associated with the β1-α1-β2 and α3 regions, which exchanged ∼10-100 times slower than the α2-β3-β4 region. This difference indicates that, of four segments that switch secondary structures, three (α2, β3, and β4) are much less stable and/or have faster opening rates than the fourth fold-switching segment (α3).

Additionally, the HDX rates for β1-α1-β2 and α3 exhibited a common and larger urea dependence than the less stable α2-β3-β4 region (**Figs. 5** and **S9**). These observations indicate that HDX for sites within β1-α1-β2 and α3 occurs through large-scale unfolding events involving unfolding of the whole protein (although the α3 helix may have some residual helical structure as HDX rates in this helix are ∼2x slower than in the β1-α1-β2 region). In contrast, the fold-switching residues across the α2-β3-β4 region undergo more frequent unfolding events.

**Figure 5.**
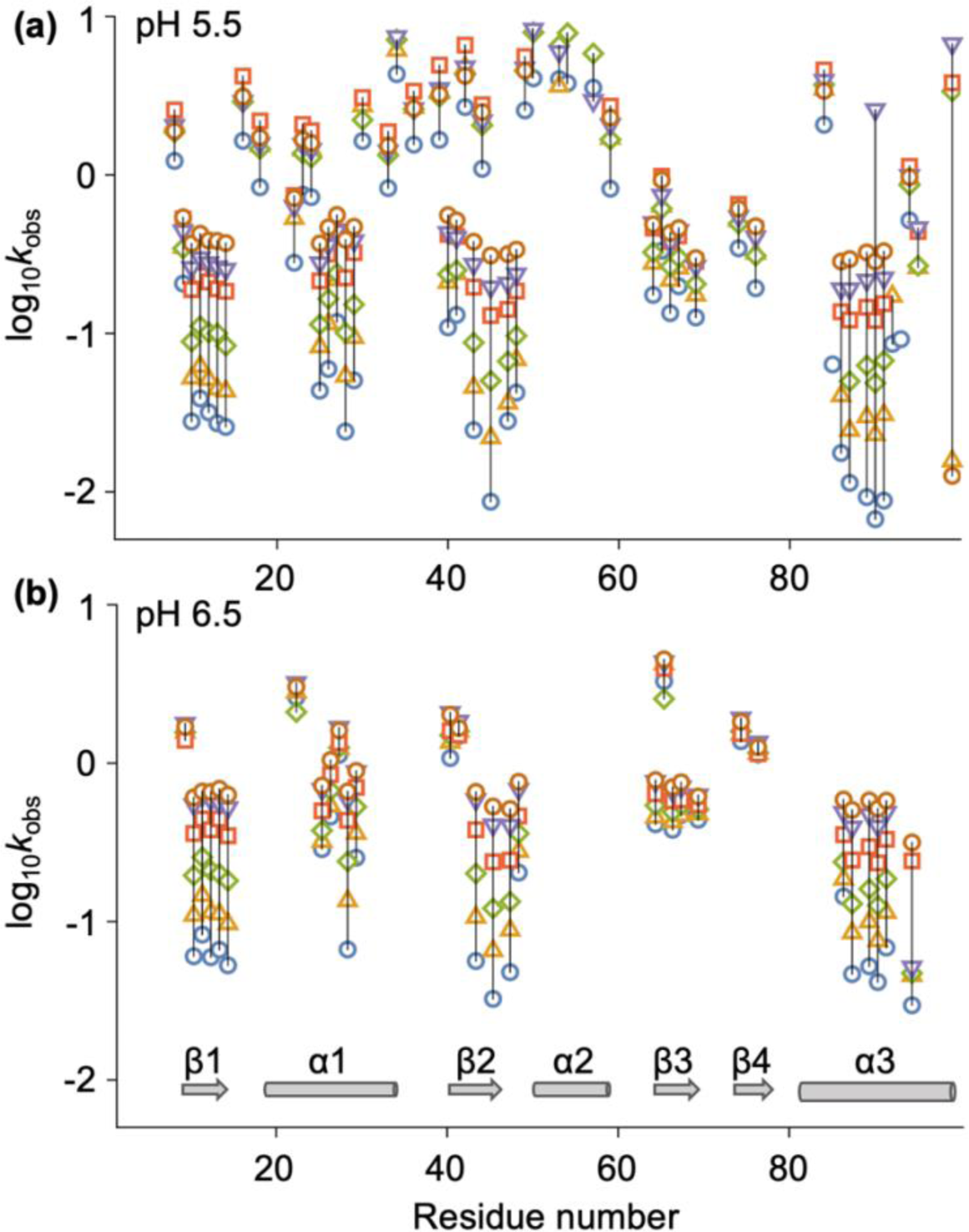
Hydrogen-deuterium exchange rates for KaiB^HDX^ indicate that β1-α1-β2 and α3 are more stable than α2-β3-β4. Observed exchange rates (log_10_ of *k*_obs_ in h^-1^) at **(a)** pH 5.5 and **(b)** pH 6.5. In **(a)** and **(b)**, blue circles, gold triangles, green diamonds, red squares, inverted purple triangles, and brown circles represent data at 0.0, 0.5, 1.0, 1.5, 2.0, and 2.5 M urea, respectively. Each data point represents the mean of two replicates, except for those at 2.0 M urea at pH 5.5 and 1.5 M urea at pH 6.5, for which there was only one measurement. The standard error in the mean (SEM) was 0.05 on average for data sets at both pH values. Residues with a measurable *k*_obs_ value at only a single urea concentration were not included in these plots.

A rigorous interpretation of HDX rates requires the determination of the extent to which exchange reflects the kinetics or thermodynamics of the opening events, termed the EX1 or EX2 limits, respectively (34). Observed HDX rates, *k*_obs_, generally depend on relative rates of opening, *k*_op_, closing (refolding), *k*_cl_, and exchange with solvent for an exposed amide proton, *k*_chem_:

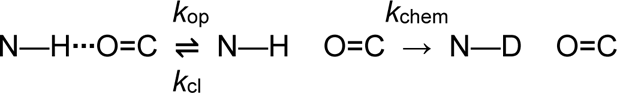

When *k*_chem_ > *k*_cl_, the system is in the EX1 limit and the HDX rate is given by *k*_obs_ = *k*_op_. Conversely, when *k*_chem_ < *k*_cl_, the system is in the EX2 limit and *k*_obs_ = *k*_chem_/PF, where PF = *K*_eq_+1 is the protection factor and *K*_eq_ = *k*_op_/*k*_cl_. As discussed in Supplementary Information, the exchange behavior for KaiB^HDX^ falls in a mixed EX1-EX2 regime, which complicates interpretation and prevents direct comparison to simulations.

Nevertheless, the patterns in **Fig. 5** described above are mirrored by site-resolved hydrogen-bond stabilities calculated from Upside simulations of ground-state dynamics of the KaiB^HDX^ construct assuming an EX2 limit (**Fig. S9**). Despite the actual exchange being in a mixed EX1-EX2 regime, we consider the HDX data consistent with the Upside model in that they both support a scenario where the α2-β3-β4 region undergoes unfolding much more frequently than the non-fold switching region, β1-α1-β2.

However, one of the fold-switching elements, the α3 helix, is much more stable than the other three and even has dynamics matching the stable, non-fold switching region. This unexpected finding argues that the energy surface does not have a stable intermediate that contains only the amino-terminal, non-fold switching half of the protein, at least in the case of KaiB^HDX^ which was engineered to be monomeric by stabilizing α3 in the fs fold at the expense of β4 in the gs fold. This finding is consistent with the significant fluxes through the globally unfolded intermediate in the Upside simulations of KaiB^D91R^. Nevertheless, both the simulations and experiments suggest there are also subglobal fold-switching pathways in which α2-β3-β4 region unfolds with the rest of the protein intact.

### Proline isomerization is the rate-determining step for KaiB^D91R^ fold switching

KaiB^D91R^ switches folds on the same hour time scale as KaiB-KaiC binding, whereas KaiB mutants locked in the fsKaiB fold by mutagenesis bind to KaiC instantaneously (5). The faster binding also abrogates clock function, suggesting that slow KaiB-KaiC binding provides an essential delay in the negative-feedback arm of the oscillator. The sensor histidine kinase, SasA, binds KaiC much faster than KaiB does because its N-terminal KaiC-binding domain is constitutively in the fsKaiB (thioredoxin-like) fold (8, 35). This much faster binding allows the SasA-KaiC complex to activate the master transcription factor, RpaA, by phosphorylation before KaiB-KaiC binding displaces SasA, creating circadian rhythms of gene expression.

Given the biological significance of the kinetics of KaiB fold switching, we wanted to identify the rate-determining step of fold switching. A key feature of the α2-β3-β4 region that more readily undergoes HDX is that it contains three prolines (P63, P70, and P72) that are *trans* in gsKaiB and *cis* in fsKaiB. Given that proline isomerization has previously been implicated in controlling protein conformational changes (36–38), and that its time scales and activation free energies (39–41) are similar to those of KaiB^D91R^ fold switching (**Fig. 3b)**, we asked whether proline isomerization is the rate-determining step for fold switching.

We found in the aforementioned Upside simulations that P63, P70, and P72 remain *trans* nearly 100% of the time in gsKaiB but are *cis* less than 30% of the time in fsKaiB. Moreover, P71, which experimentally is *trans* in both gsKaiB and fsKaiB, populates the *cis* isomer 90% of the time in simulations of fsKaiB (**Table S2**). These deviations from experimental observations likely result from Upside representing the proline *cis* and *trans* states through a double-well potential without accounting for side chain steric hindrance, so we interpret P70-P72 as acting as a single unit in the Upside model of KaiB^D91R^.

With this limitation of our model in mind, we probe the structural changes associated with proline isomerization. The *cis* fraction of P63 and any of P70–P72 rises sharply when *Q*_fs,C_ – *Q*_gs,C_ ≈ 0.5 (**Fig. S10a, b**), near the transition state for fold switching (**Fig. S6**). However, the P70–P72 isomerization appears less temperature sensitive and less dependent on the *Q*_N_ value, which suggests that it does not require global unfolding. Rather, consistent with the location of P70–P72 in the sequence, their isomerization occurs with the packing of the two C-terminal β-strands, β3 and β4, in the fsKaiB state (**Fig. S10c**).

To probe whether isomerization of P63, P70, and P72 (with P71 in the *trans* state) is sufficient to induce fold switching computationally, we altered the conformations of P63, P70, and P72 in the fsKaiB structure from *cis* to *trans* and ran simulations of KaiB^D91R^ with each proline locked in its assigned conformer (i.e., subjected to a single-well potential for backbone isomerization). From 240 such simulations (the temperature was varied to 0.89 or 0.90 relative to a melting temperature of 0.94), we observed four transitions to gsKaiB, all of which involved subglobal (and not global) unfolding (**Figs. 6a** and **S11**). In these simulations, the transition tends to start with β3_fs_ and β4_fs_, which may give the proline(s) the freedom to isomerize; the remaining steps involve α3_fs_ unfolding followed by folding of α3_gs_ and either β4_gs_ or α2_gs_. 384 analogous simulations starting in the gsKaiB state with P63, P70, and P72 locked in *cis* did not yield fold switching to the fsKaiB state for KaiB^D91R^. This difference could reflect the relative stability of the gsKaiB state for the KaiB^D91R^ mutant compared to the fsKaiB state (by contrast, KaiB^G89A^ displayed both gs→fs and gs←fs transitions during analogous Upside simulations (42)). Nonetheless, our observation of proline isomerization-induced, subglobal gs←fs fold-switching events indicates that global unfolding is not required for fold switching, despite the global unfolding in the unbiased simulations reported above.

**Figure 6.**
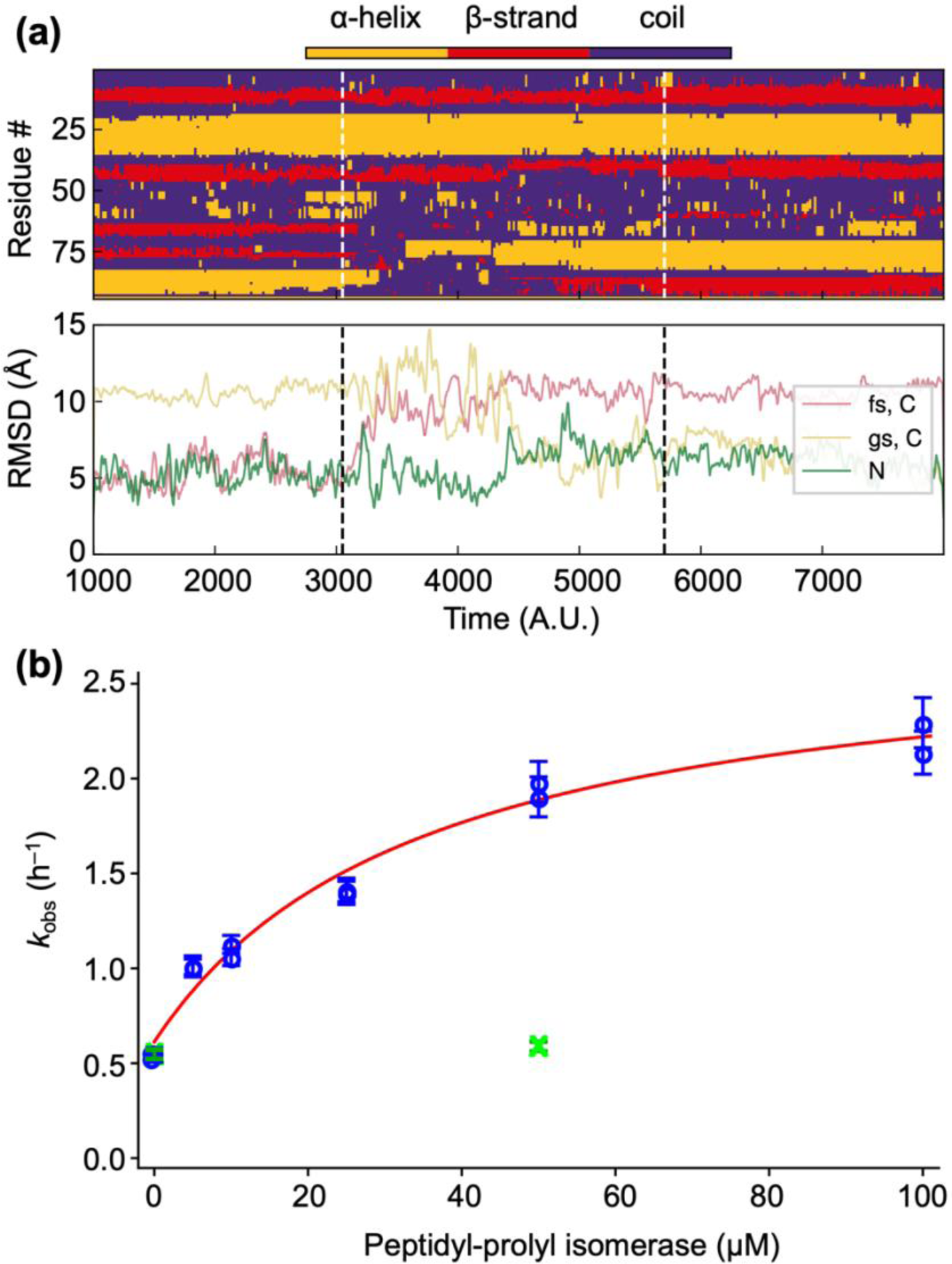
Proline isomerization is rate-limiting for fsKaiB→gsKaiB fold-switching of KaiB^D91R^. **(a)** An Upside simulation starting from fsKaiB with P63, P70, and P72 fixed in *trans*. (top) Secondary structure assigned via DSSP (79) and (bottom) RMSD to either the C-terminal half of fsKaiB, the C-terminal half of gsKaiB, or the N-terminal half (**Table S1**) during a section of a successful fold-switching simulation from fsKaiB to gsKaiB. RMSDs are plotted as a moving average over 20 frames. The dotted lines indicate the approximate start and end times of the fold-switching event. **(b)** Observed rates of fold switching, *k*_obs_, after a jump to 35 °C for KaiB^D91R^ samples pre-equilibrated at 4 °C as a function of the concentration of human peptidyl prolyl isomerase A (PPIA). 190–200 µM ^15^N-enriched KaiB^D91R^ samples stored at −80 °C were incubated at 4 °C for 24 h before inserting them into the NMR spectrometer, which was set to a sample temperature of 35 °C. Twenty-three ^15^N,^1^H HSQC spectra were collected every 18 min over a span of seven hours. Residue-specific rates were determined from these spectra as described in Supplementary Information. Open blue circles represent the means of *k*_obs_ values for residues with resolved and assigned gsKaiB and fsKaiB peaks in the absence of the PPIA inhibitor, cyclosporin A (CsA). The green symbols represent means of *k*_obs_ values where 55 µM CsA was added to the samples. Error bars represent the standard deviation of the rates about the mean. Error bars are not shown when they are smaller than the size of the symbol.

To verify the computational prediction that proline isomerization is a key step in fold switching, we modulated the barrier to proline isomerization using human peptidyl prolyl isomerase A (PPIA, also known as cyclophilin A or CypA (43)), which is a member of the cyclophilin family. PPIA enhanced the rate of KaiB fold switching in a dose-dependent manner as measured by NMR (**Fig. 6b**), and cyclosporin A, an inhibitor of cyclophilins (43), abolished the effect of PPIA, suggesting that proline isomerization indeed dictates the rate of fold switching. Given that prolyl isomerases such as PPIA likely bind to prolines in unstructured segments (37, 44, 45), unfolding of β3_fs_ and β4_fs_ likely precedes proline isomerization, consistent with the simulations. Our simulations and experiments do not resolve whether any of the three isomerizing proline residues plays a dominant role. Nevertheless, they suggest that it is the combination of unfolding and proline isomerization that makes KaiB fold switching slow.

## DISCUSSION

Over the last couple decades, there has been a dramatic breakdown in the traditional one-to-one-to-one relationship between sequence, fold, and function of proteins. Not only are there intrinsically disordered proteins (46), some of which fold upon binding, but also amyloid proteins and prions, which can change their folds but do so irreversibly (47). Metamorphic proteins like KaiB, which can populate two well-folded structures reversibly, have received much less attention. They are distinct from morpheeins, which are proteins that adopt different quaternary structures while retaining the same fold (48), and moonlighting proteins, which have more than one biochemical function but the same fold (49). It remains to determine how many metamorphic proteins in the Protein Data Bank are masquerading as single-fold (or monomorphic) proteins (2, 50–52). Our work aids in this effort by providing insights into the forces that mediate fold switching.

Our simulations and native-state HDX data on KaiB^HDX^ suggest that KaiB switches folds via both subglobal and global pathways, with the α2-β3-β4 region of fsKaiB exhibiting the greatest lability. These mechanistic findings can be linked to our temperature studies through the observation that electron-spin resonance experiments on *T. elongatus* KaiB labeled at different positions with paramagnetic probes indicate that internal motions in the α2 region of the gsKaiB fold are particularly sensitive to temperature (53). These residues are at a monomer-monomer interface in the WT KaiB homotetramer. Given that oligomerization requires KaiB to be in the gsKaiB fold (10, 11), our finding that an increase in temperature stabilizes gsKaiB relative to fsKaiB may explain the observation that a mutant of *T. elongatus* KaiB with the first nine residues deleted is a dimer at 35 °C but a monomer at 4 °C (54); we similarly interpret the observation that lower temperature favors formation of the KaiABC complex (55). Whether oligomerization is rate limiting for WT KaiB, as suggested by a computed thermodynamic cycle and structure-based modeling (14), remains to be determined because our finding that proline isomerization is rate limiting for fold switching was obtained for a monomeric construct (KaiB^D91R^). However, we note that slow conformational switching in the protein BMAL1, which plays significant roles in timekeeping by the mammalian circadian clock, requires proline isomerization and is modulated by isomerases of the cyclophilin family (36).

In the Protein Data Bank, the gsKaiB fold is found only in KaiB homologs, whereas the fsKaiB thioredoxin-like fold is common. Thus, it is plausible that ancestors of KaiB adopted the thioredoxin-like fold exclusively and later evolved the ability to reversibly switch to the gsKaiB fold for the purpose of circadian timekeeping. For example, the Kai system in *Rhodobacter sphaeroides* functions as an hourglass timer rather than a self-sustaining clock, and its KaiB can adopt both the gsKaiB and fsKaiB states (50, 56). In contrast, the bacterium *Legionella pneumophila*, which is not known to possess a circadian clock, has a *kaiBC* operon where the KaiB protein crystallizes in the thioredoxin-like fsKaiB fold and has an alanine residue at position 89 as KaiB^G89A^ does (57).

By controlling KaiBC complex formation (5, 17, 58), KaiB fold switching *(i)* contributes to the slowness of the circadian clock, *(ii)* provides time-delayed negative feedback essential to all biochemical oscillators (59), and *(iii)* opens a transient window of RpaA activation by allowing SasA to bind to the CI domains of KaiC before KaiB does (8). KaiB assembles as a hexameric ring onto the post-ATP hydrolysis state of the CI ring at night (13, 55, 56), displaces SasA (15), sequesters an autoinhibited state of KaiA (13, 55), and separately binds CikA, which then dephosphorylates RpaA. When bound to KaiC, KaiB also interacts with the protein KidA to tune the period of the cyanobacterial circadian oscillator (19). Thus, fold switching is essential to multiple aspects of clock function.

A remarkable feature of circadian clocks is that their amplitudes change so that their periodicities are relatively insensitive to temperature (60–63). This temperature compensation enables circadian clocks to reliably maintain an internal representation of time. For cyanobacteria, it is well established that the ATPase activity of KaiC is temperature compensated (62, 64). The temperature shifts that we observe for the gsKaiB ⇌ fsKaiB equilibrium should also contribute to temperature compensation of the cyanobacterial circadian clock. To illustrate this, we used a simple reaction scheme to model the binding kinetics between KaiB^D91R^ and the CI domain of KaiC (**Fig. 7a**). This model can be described by the following coupled differential equations (see Supplementary Information for details):

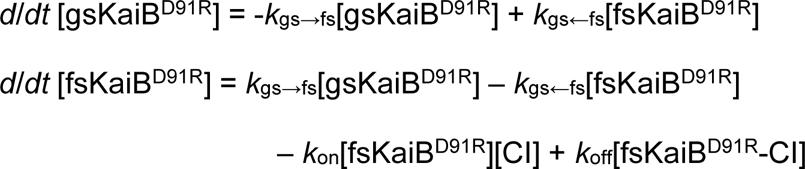

**Figure 7.**
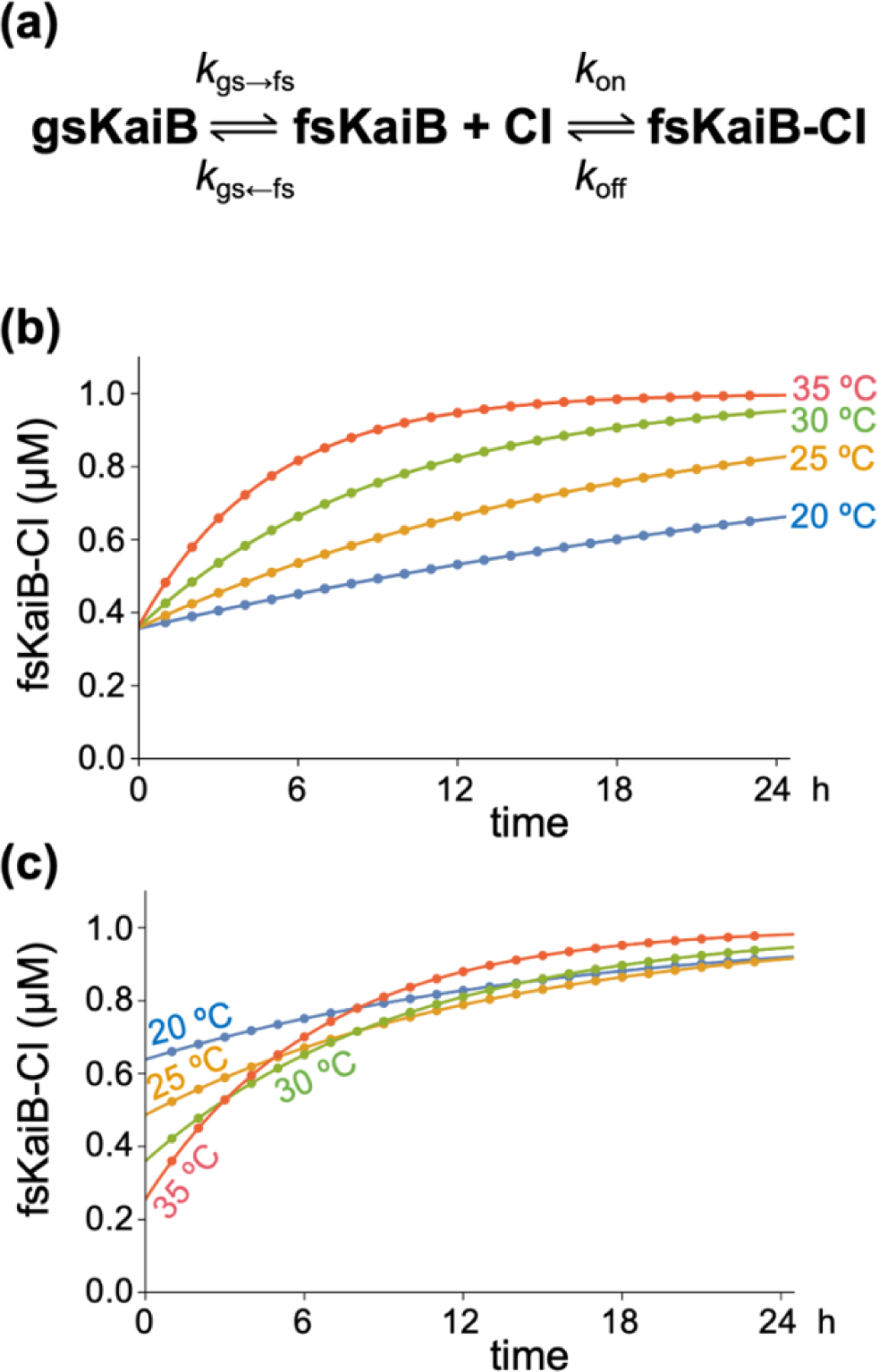
Kinetics of formation of the fsKaiB^D91R^-CI complex is partially temperature compensated. **(a)** Reaction scheme of KaiB^D91R^ fold switching and binding the CI domain of KaiC. **(b)** Calculated kinetics of fsKaiB^D91R^-CI complex formation when the thermodynamics and kinetics are fixed at their experimentally determined 30 °C values. **(c)** Calculated kinetics of fsKaiB^D91R^-CI complex formation using thermodynamics and kinetics values experimentally determined at each temperature modeled.

In **Fig. 7b**, we calculated the kinetics of fsKaiB^D91R^-CI complex formation as a function of temperature but with Δ*G*, *k*_gs→fs_, and *k*_gs←fs_ fixed at values experimentally determined at 30 °C. In contrast, for **Fig. 7c** we used values determined experimentally for each temperature modeled. As can be seen for both cases, higher temperatures lead to faster binding. However, for the temperature-dependent *k*_gs→fs_ and *k*_gs←fs_ case, the amounts of fsKaiB^D91R^-CI complex formed across the different temperatures are more similar, because with increasing temperature the gsKaiB^D91R^ ⇌ fsKaiB^D91R^ equilibrium shifts toward gsKaiB^D91R^, and Δ*G*^‡^_gs→fs_ and Δ*G*^‡^_gs←fs_ increase and decrease, respectively. Thus, KaiB fold switching likely contributes to the overall mechanism of temperature compensation in the cyanobacterial circadian clock. Interestingly, the cyanobacterial circadian clock can also compensate for changes in molecular crowding (65), which not only impacts complex formation but also shifts the gsKaiB ⇌ fsKaiB reaction to the left (66). Mechanistic studies of the interplay between clock complex formation and fold switching under various conditions would be worthwhile.

## MATERIALS & METHODS

### Cloning, Protein Expression, and Purification

KaiB constructs are listed in **Table S3** and were expressed and purified using previously published methods (67, 68). Briefly, cells were grown to OD_600_ ≈ 0.6 in M9 medium containing ^15^N-labeled ammonium chloride (Cambridge Isotope Laboratories, Inc.) and induced by adding isopropyl β-d-1-thiogalactopyranoside to a final concentration of 200 µM for 12 h at 30 °C. Cells were harvested and cell pellets were resuspended using 4 °C lysis buffer (50mM NaH_2_PO_4_, 500mM NaCl, pH 8.0) and homogenized with an Avestin C3 Emulsiflex homogenizer (Avestin Inc, Canada). Supernatant was clarified by centrifugation, loaded onto a Ni-NTA column (QIAGEN, #30230), and the column was washed with a buffer solution (50 mM NaH_2_PO_4_, 500 mM NaCl, 20 mM imidazole, pH 8.0). His-tagged KaiB was eluted with a buffer solution consisting of 50 mM NaH_2_PO_4_, 500 mM NaCl, and 250 mM imidazole, pH 8.0. His-tagged ULP1 protease was added to His-tagged KaiB to final concentration of 3 µM and the sample was incubated for 12 h followed by loading 10x-diluted sample onto a Ni-NTA column to remove ULP1 and the SUMO tag. The flowthrough containing KaiB was then passed through a HiLoad 16/600 Superdex 75 size-exclusion column in phosphate buffer (20 mM Na_2_HPO_4_/NaH_2_PO_4_, 100 mM NaCl, pH 7.0) by FPLC. Finally, protein purity was assessed by SDS PAGE. All KaiB samples were stored at −80 °C.

### Analytical Gel-Filtration Chromatography

A Superdex™ 75 Increase 10/300 GL column (Cytiva) with an injection volume of 100 µL was used for analytical gel-filtration chromatography of KaiB samples in SEC buffer (20 mM Na_2_HPO_4_/NaH_2_PO_4_, 100 mM NaCl, pH 7.0). Samples were incubated at 4 °C overnight (∼12 h) and then injected onto the column with a flow rate of 0.7 mL/min.

### NMR Temperature-Jump Experiments

NMR experiments were performed on a Bruker Avance III 600 MHz spectrometer equipped with a TCI cryoprobe. NMR samples contained 200 μM ^15^N-enriched *T. elongatus* KaiB 1-94 Y8A D91R Y94A (KaiB^D91R^) in 350 μL NMR buffer (20 mM Na_2_HPO_4_/NaH_2_PO_4_, 100 mM NaCl, 0.02% NaN_3_,10 μM DSS, pH 7.0, 90% H_2_O, 10% D_2_O). KaiB^D91R^ samples were stored at −80 °C and incubated at 4 °C for 24 h prior to NMR experiments. Samples were not re-used. A series of two-dimensional ^1^H-^15^N HSQC spectra were recorded at 293 K, 298 K, 303 K and 308 K, respectively. ^1^H chemical shifts were referenced to internal DSS and ^15^N chemical shifts were indirectly referenced to DSS using absolute frequency ratios listed on the BMRB website. NMR data were processed with NMRPipe and analyzed using nmrDraw or NMRFAM-Sparky (69, 70). See Supplementary Information for details on data analysis.

### NMR Hydrogen-Deuterium Exchange (HDX) Experiments

HDX experiments were initiated by adding 12 °C D_2_O to lyophilized ^15^N-enriched *T. elongatus* KaiB 1-99 Y8A P71A G89A D91R Y94A (KaiB^HDX^) powder followed by acquisition of ^15^N,^1^H HSQC spectra over a 23-h period at a sample temperature of 12 °C and urea concentrations of 0, 0.5, 1.0, 1.5, 2.0, and 2.5 M urea. Prior to lyophilization, samples were either at pH 5.5 or pH 6.5. HDX rates were determined by fitting HSQC peak heights as a function of time. Under base-catalyzed exchange (pH > 3.5), urea can interfere with HDX and thus we corrected our measured rates accordingly (71). HSQC peaks of slowly exchanging residues can transiently increase in intensity if nearby residues undergo much more rapid HDX (72). Thus, we fit individual HSQC peak intensities, *I*(*t*), to the following equation:

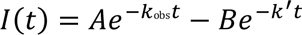

where the first term accounts for HDX of the residue in question and the second term accounts for the effect of line narrowing produced by rapid deuteration of nearby residues.

### Upside Simulations

We used a near-atomic resolution model called Upside to elucidate fold-switching pathways and their statistics. In Upside simulations, protein conformations are represented by explicit N, C_α_, and C atoms, but energies are calculated so as to account for the expected positions of the amide proton, carbonyl oxygen, and an oriented side-chain bead (effectively 6 atoms per residue) (26, 73, 74). The energy function includes hydrogen bonds, side chain-side chain and side chain-backbone interactions, and desolvation terms, along with trained neighbor- and residue-dependent (ɸ, ѱ) dihedral maps. The force field parameters were learned using the contrastive divergence method.

Upside evolves atom positions with Langevin dynamics. A unique feature of the model is that it represents the side chain packing probabilistically. As a consequence, side chain packing instantly equilibrates to the backbone conformation at every molecular dynamics (MD) step, which makes each step equivalent to about 30 ps of all-atom dynamics. These elements of the Upside mode allow it to describe protein dynamics accurately and rapidly with considerable molecular detail while still generating thermally equilibrated ensembles.

Despite Upside’s speed, prohibitively long times are required to observe fold switching in unbiased MD simulations. However, we can obtain statistical information about fold-switching pathways through an approach called the dynamical Galerkin approximation (DGA) (27, 28, 75, 76). In this approach, relatively short unbiased simulations are used to sample portions of the fold-switching pathways, and these data are then used to estimate averages that determine the coefficients of basis expansions of functions of the dynamics, subject to a Markov assumption. Importantly, DGA does not require any single simulation to connect gsKaiB and fsKaiB but simply that the pathways between them are well sampled in aggregate by the short simulations.

In Supplementary Information, we describe the parameterization of the Upside proline potential, calibration of the Upside temperature, initialization of the Upside simulations, collective variables that we use to characterize the dynamics, details of the DGA calculations, and the Upside HDX analysis.

### All-atom simulations

We carried out all-atom MD simulations to determine the temperature dependence of the relative free energies of the gsKaiB and fsKaiB conformations. At each temperature from 280 K to 350 K in steps of 5 K, we simulated gsKaiB and fsKaiB for 2 μs and collected the last 1 μs for analysis by MBAR (77). See Supplementary Information for additional details.

## ACKNOWLEDGEMENTS

This work was supported by US National Institutes of Health (NIH) grants R35GM144110 (to A.L.), R35GM136381 (to A.R.D.), R35GM148233 (to T.R.S.) and R35GM141849 (to C.L.P), National Science Foundation (NSF) MCB grants 1953402 (to A.R.D.) and 2023077 (to T.R.S.), NSF-CREST: Center for Cellular and Biomolecular Machines at the University of California, Merced (NSF-HRD-1547848), and US Army grant W911NF-23-1-0248 (to A.L.). S.C.G. was supported by a NSF Graduate Research Fellowship under Grant No. 2140001. We also acknowledge the Health Sciences Research Institute at UC Merced for pre- and post-award management. We thank Yong-Gang Chang, Roger Tseng, Madhurima Das, and Shahar Sukenik for valuable advice, David Rice for NMR support, and Nabil Faruk, Nick Bayhi, and Darren Liu for Upside-related discussions and scripts. This work was completed in part with resources provided by the University of Chicago Research Computing Center and we are grateful for their assistance with the calculations. The “Beagle-3: A Shared GPU Cluster for Biomolecular Sciences” is supported by the NIH under the High-End Instrumentation (HEI) grant program award 1S10OD028655-0.

