## Supplementary Information for "Temperature-Dependent Fold-Switching Mechanism of the Circadian Clock Protein KaiB"

*Co-first authors

[#]Co-corresponding authors

### SUPPLEMENTARY TEXT

#### EXPERIMENTAL DETAILS

##### Measuring residue-specific Δ*G* values for gsKaiB$^{D91R}$ ⇌ fsKaiB$^{D91R}$.

Δ*G* values for the gsKaiB$^{D91R}$ ⇌ fsKaiB$^{D91R}$ reaction can be estimated from integrated HSQC peak volumes on equilibrated samples as follows:

$$\Delta G = -RT\,\ln K_{eq}$$

with

$$K_{eq} = p_{fs,eq}/p_{gs,eq},$$

$$p_{gs,eq} = V_{gs,eq}/(V_{gs,eq} + V_{fs,eq}),$$

and

$$p_{fs,eq} = V_{fs,eq}/(V_{gs,eq} + V_{fs,eq}),$$

where, $p_{gs,eq}$ and $p_{fs,eq}$ are the residue-specific fractional equilibrium populations of gsKaiB$^{D91R}$ and fsKaiB$^{D91R}$, respectively, and $V_{gs,eq}$ and $V_{fs,eq}$ are the corresponding HSQC peak volumes of gsKaiB$^{D91R}$ and fsKaiB$^{D91R}$ under equilibrium conditions.

##### Determination of residue-specific $\Delta G^{‡}_{gs\rightarrow fs}$ and $\Delta G^{‡}_{gs\leftarrow fs}$ values from temperature-jump experiments on KaiB$^{D91R}$ (and KaiB$^{G89A}$).

The observed rates of $^{15}$N-enriched KaiB$^{D91R}$ fold switching, $k_{obs}$, upon a jump in sample temperature were obtained by fitting changes in fractional populations, $p_{gs}$ and $p_{fs}$, determined from volume integration of HSQC peaks to the following equations:

$$p_{gs} = 1 - (\mathrm{A}\, e^{-k_{obs}\, t} + \mathrm{B})$$

$$p_{fs} = \mathrm{A}\, e^{-k_{obs}\, t} + \mathrm{B}$$

Residue-specific forward and reverse rate constants, $k_{gs\rightarrow fs}$ and $k_{gs\leftarrow fs}$, of the gsKaiB$^{D91R}$ ⇌ fsKaiB$^{D91R}$ reaction were determined for each temperature as follows:

$$k_{gs\rightarrow fs} = k_{obs}\ p_{fs,eq}$$

$$k_{gs\leftarrow fs} = k_{obs}\ p_{gs,eq}$$

These rate constants allowed us to determine the activation free energies, $\Delta G^{\ddagger}_{gs\rightarrow fs}$ and $\Delta G^{\ddagger}_{gs\leftarrow fs}$, using transition-state theory:

$$\Delta G^{\ddagger}_{gs\rightarrow fs} = -RT\ \ln(h\ k_{gs\rightarrow fs}\ /k_B T)$$

$$\Delta G^{\ddagger}_{gs\leftarrow fs} = -RT\ \ln(h\ k_{gs\leftarrow fs}\ /k_B T)$$

**The exchange behavior of KaiB$^{HDX}$ is in a mixed EX1-EX2 regime.**

One method to identify the extent to which a residue is in the EX1 or EX2 regime is to examine its HDX rate as a function of pH, taking advantage of the known linear dependence of log($k_{chem}$) on pH. Since $k_{obs}$ is independent of $k_{chem}$ under EX1 conditions, $k_{obs}$ will not be affected by pH and hence, the plot of log($k_{obs}$) versus pH will have zero slope, provided the stability of the protein does not change over the chosen pH range. In contrast, the slope should be one under EX2 conditions where $k_{obs}$ is proportional to $k_{chem}$. An assumption underlying this approach is that the stability of the protein does not change over the pH values tested; we confirmed that this is the case by thermal melting studies tracking far-UV circular dichroism for KaiB$^{HDX}$ (**Fig. S12**).

We observed that the slopes in the $\log_{10}(k_{obs})$ versus pH plot for most sites are between 0.2 and 1. Because the slopes in the EX1 and EX2 limits are 0 and 1, respectively, the exchange behavior falls in a mixed EX1-EX2 regime and is dependent on the particular residue (**Fig. S13**). That said, we expect that the exchange is closer to the EX1 limit. In addition, elevated pH and urea concentrations amplify the probability of the hydrogen-deuterium exchange (HDX) data being in the EX1 regime because these two factors increase the intrinsic exchange rate and decrease the refolding rate, respectively, thereby increasing the likelihood that they satisfy the EX1 condition $k_{chem} > k_{cl}$. Specifically, the chemical exchange rate ($k_{chem}$) is accelerated 10-fold at pH 6.5 versus pH 5.5, while the folding rate ($k_{cl}$) is reduced at higher urea concentrations. In the EX1 limit, $k_{obs} = k_{unf}$ (i.e., $k_{op}$), and hence $k_{obs}$ should be independent of $k_{chem}$, whereas in the EX2 limit, $k_{obs} = k_{chem}$/PF, and thus dependent on $k_{chem}$. These behaviors, along with $k_{chem}$'s inherent increase by 10x for every pH unit, imply that the slope of the $\log_{10}(k_{obs}/k_{chem})$ plot in the EX1 limit should be negative but zero in the EX2 limit. Since the observed slopes are predominantly negative, the data better support exchange being in the EX1 limit. Overall, the pH dependence of the rates suggests that the exchange for these sites is closer to the EX1 limit, especially with added denaturant. This in turn implies that the $k_{obs}$ values mostly reflect the unfolding rate of a large-scale opening event.

#### COMPUTATIONAL DETAILS

##### Systems studied with Upside.

We used CHARMM-GUI version 3.7 (1) to prepare systems for Upside simulations. The X-ray crystal structure for tetrameric wildtype (WT) ground state KaiB was retrieved from the Protein Data Bank (PDB ID 2QKE (2)). Monomeric gsKaiB was isolated by modeling only chain B, since all its residues were resolved. The NMR structure of a fold-switched KaiB mutant was also retrieved from the Protein Data Bank (PDB ID 5JYT (3)), and CHARMM-GUI was used to mutate the sequence back to the WT sequence. Both gsKaiB and fsKaiB structures were truncated and mutated to the $KaiB^{D91R}$ sequence or the $KaiB^{HDX}$ sequence (**Table S3**). All simulations were performed and analyzed with a combination of Upside (4, 5) and MDTraj 1.9.9 (6). All structures were visualized in VMD (7).

**Parameterization of Upside proline isomerization potential.**

To allow for the explicit isomerization of the prolines during the unbiased dynamical Galerkin approximation (DGA) simulations, we employed a two-well proline isomerization potential. The minima are at $\omega$ dihedral angle values of 180° (*trans*) and 0° (*cis*) and are separated by a 2.5–5.0 $k_BT$ barrier. This barrier is set much lower than the experimental value of 76-92 kJ/mol (8–10) to enable adequate sampling of the proline isomerization states during unbiased simulations. Despite this choice, we expect to obtain meaningful pathways because we adjust the difference in energy between the *cis* and *trans* energy minima ($\Delta E_{cis/trans} = E_{cis} - E_{trans}$) to match literature values (11, 12). To set $\Delta E_{cis/trans}$, we ran a series of simulations of five-amino acid fragments, each with at least one proline that was allowed to isomerize. The literature values for $\Delta E_{cis/trans}$ are dependent on the preceding amino acid (11, 12); in KaiB, the seven prolines are preceded by one of four unique amino acids (Xaa = Ala, Leu, Thr, Pro). For all small peptide simulations, a five amino acid segment was taken from PDB ID 5JYT and mutated and capped using CHARMM-GUI to the sequence $H_3CCO$-Ala-Xaa-Pro-Ala-Lys-$NH_2$. For each peptide, we ran eight independent unbiased simulations of length 20 million Upside time units initialized from each isomerization state (*cis* or *trans*), where

$\Delta E_{\text{cis/trans}}$ ranged from –1.0 to 2.7 $k_B T$. The proline dihedral angle was calculated, and the proline was considered *cis* if $\omega \in [-90°, 90°]$.

**Upside temperature calibration.**

To investigate the role of temperature in the fold-switching dynamics of KaiB, we needed to set a simulation temperature, as the Upside temperature scale is uncertain to within 10–20º (13). We ran 8 sets of temperature replica-exchange (T-REMD) simulations of $\text{KaiB}^{\text{D91R}}$, each with 48 replicas. We initialized 8 sets of T-REMD simulations from the gsKaiB, fsKaiB, and intermediate structures characterized by the secondary structure collective variables (**Collective variables**). Prolines were allowed to isomerize under a double-well potential with a barrier of 2.5 $k_B T$ (**Parameterization of proline isomerization potential**), to allow more rapid equilibration of the *cis*/*trans* states. Each simulation was 3 million Upside time units long at temperatures ranging between $T$ = 0.78 and $T$ = 1.02 (in units of $k_B$), with exchanges attempted every 10 time units. Coordinates were saved every 50 time units. The first 500,000 frames of each trajectory were discarded as burn-in samples. After reweighting all 384 trajectories with the multistate Bennett acceptance ratio (MBAR) method (14), we determined the melting temperature of fsKaiB to be $T$ = 0.92 (**Fig. S14**), the point where the number of KaiB hydrogen bonds as a function of simulation temperature decreased and the radius of gyration increased dramatically.

**Initialization of Upside simulations.**

To achieve good coverage of the fold-switching pathways for the DGA simulations, we did the following: as described in **Upside temperature calibration**, we performed 8 x 48 simulations using T-REMD to adequately sample the conformational space of gsKaiB, fsKaiB, and unfolded intermediates. We drew 32 structures evenly spaced by 1000 frames (50,000 time steps) starting from the 10,000$^{th}$ frame from each T-REMD trajectory. From each of the 8 x 48 x 32 starting structures, we launched 2 independent unbiased simulations of length 200,000 time units, one initialized with P63, P70, P71, and P72 in the *cis* state, and one initialized with the prolines in the *trans* state. In these simulations, all prolines were subject to a double-well potential with a barrier of 5 $k_B T$. We generated data sets in this way at Upside temperatures of 0.87, 0.89, 0.91, which are near but below the melting temperature (see the **Upside temperature calibration**). We saved configurations every 200 time units.

**Collective variables.**

To characterize the structures in Upside simulations, we define a number of collective variables (CVs) based on differences in fractions of contacts in the gsKaiB and fsKaiB states. We compute the fraction of contacts between non-hydrogen atoms using the formula in ref. (15):

$$Q_s = (1/N)\sum_{ij}\{1+\exp[\beta(r_{ij} - \lambda s_{ij}^0)]\}^{-1},$$

where the sum runs over non-hydrogen atoms within 0.45 nm in reference structure *s* (*s* = gsKaiB or fsKaiB as constructed as described in **Systems studied with Upside**), $r_{ij}$ is the distance in the structure of interest, $s_{ij}$ is the distance in reference structure $s$, and $N$ is a normalization factor; only $ij$ pairs from residues separated by three or more sequence positions were included. We set the β and λ parameters to be 50 nm$^{-1}$ and 1.8, respectively. β determines the "softness" of the switching function, determining how quickly the contact function decays from 1 to 0 after the reference distance is passed. λ is the tolerance for the reference distance, providing slack to allow atoms at near-native distance to still be considered contacts.

As noted in the main text, we also considered subsets of the contacts in the gsKaiB and fsKaiB states. These included fractions of contacts for the N-terminal half and the C-terminal half denoted $Q_{\mathrm{N}}$, $Q_{\mathrm{gs/fs,C}}$. We also considered fractions of contacts for individual fold-switching secondary structure elements, following a similar naming convention. While the definition above accurately captured the melting/repositioning of α-helices, we found it too sensitive to register shifts of β-sheets. Thus, for β-strands, we modify the definition of $Q_s$ and instead measure the number of contacts to neighboring β-strands, normalized to the number of contacts between those strands in the experimental structures. For $Q_{\mathrm{fs},\beta3}$ we measured contacts between β3 and β1 in fsKaiB. For $Q_{\mathrm{fs},\beta4}$ we measured contacts between β4 and β3 in fsKaiB. For $Q_{\mathrm{gs},\beta4}$ we measured contacts between β4 and β1 in gsKaiB. See **Table S1** for definitions of the secondary structures and **Fig. 1** for their arrangements. Because it is possible for non-native conformations to have more overall contacts between specified β-strands than the experimental structure does, $Q$ for β-strands ranges from 0 to approximately 1.3. This means that the typical ranges for $Q_{\mathrm{fs},\alpha2} - Q_{\mathrm{gs},\beta3}$ and $Q_{\mathrm{fs},\alpha3} - Q_{\mathrm{gs},\beta4}$ CVs are between –

1.3 and +1, while the typical ranges for $Q_{fs,\beta3} - Q_{gs,\alpha2}$ and $Q_{fs,\beta4} - Q_{gs,\alpha3}$ are between –1 and +1.3.

**DGA weights and basis set construction.**

Because the unbiased trajectories used to compute kinetic statistics in DGA are not sampled from a stationary, equilibrium distribution in general, typically one must reweight the sampled data to the correct equilibrium distribution. Mathematically, given a sampling distribution μ that we want to reweight to an equilibrium distribution π we have for any function *f*

$$\langle f\rangle_\pi = \langle wf\rangle_\mu/\langle w\rangle_\mu$$

where $w = \pi/\mu$ are the weights and bracket subscripts indicate the distributions used for the averages. Typical approaches for computing the weights (16, 17) yielded poor agreement with the equilibrium distribution sampled with T-REMD, which we expect to be reasonably accurate. Therefore, we instead chose to set the weight of each frame in our unbiased dataset to the MBAR weight of the initial frame in the trajectory. This assumes that the MBAR weights are stationary over the trajectory. This is not strictly true because the T-REMD simulations are of finite duration and we initialize isomerizing prolines with equal weights even though we expect their weights to be different at equilibrium. In practice, we found this procedure yielded potentials of mean force from DGA (**Fig. 4**) in good agreement with the T-REMD/MBAR results (**Fig. S15**).

To generate the basis for computing the probability of reaching the fsKaiB state before the gsKaiB state, we first used PyEMMA (18) to compute molecular features from our existing unbiased trajectory data. We computed contacts between residue pairs more than three residues away in the set 3, 9, 11, 13, 16, 23, 33, 38, 41, 43, 45, 47, 51, 56, 59, 63, 65, 67, 70, 72, 75, 78, 81, 84, 87, 90, and 93. Contacts were counted when two residues had any heavy-atom distance between them less than 0.45 nm. We then reduced the dimensionality of the binary contact maps to five dimensions using the integrated variational approach to conformational dynamics (IVAC) (19), integrating over lag times from 1 to 200 frames (200 to 40,000 time steps). We combined the five IVAC coordinates with cos($\omega$) for P63, P70, P71, and P72, to emphasize the importance of the proline isomerization state, and then clustered the resulting nine-dimensional data into 500 clusters using mini-batch *k*-means in scikit-learn (20). Each cluster defined a basis function that was one on the cluster and zero otherwise. The results were only mildly sensitive to the size of the basis set (the number of clusters).

To obey the homogeneous boundary conditions of the DGA operator equations, we defined our gsKaiB and fsKaiB states in terms of our previously described CVs using the potentials of mean force from T-REMD and distributions of CVs from unbiased simulations. The cutoffs for the CVs we used are listed in **Table S4**. For calculating the probability of reaching the fsKaiB state before the gsKaiB state ($q$), we used a function that was one in the fsKaiB state and zero elsewhere as the DGA "guess" function that accounts for the boundary conditions (16, 21), and we used an improved algorithm that incorporated memory (22) with a lag time of 100 frames (20,000 time steps) and one memory kernel.

**Reactive flux through unfolded state.**

To separate reactive trajectories (that is, trajectories which fold-switch) that pass through a globally unfolded (i.e., N-terminal half unfolded) state from all reactive trajectories, we follow the procedure from ref. (23), which treats reactive trajectories in open domains. Following their notation, we are interested in dynamics of a Markov process $X_t$ in some domain $S$ comprised of a "open" domain $O \subset S$ and an artificial state $\omega$ that closes the dynamics. Here, we define the unfolded state $\omega$ as containing structures where $Q_{\mathrm{N}} < 0.16$, $|Q_{\mathrm{fs},\beta3} - Q_{\mathrm{gs},\alpha2}| < 0.5$, $|Q_{\mathrm{fs},\beta4} - Q_{\mathrm{gs},\alpha3}| < 0.5$, and $|Q_{\mathrm{fs},\alpha3} - Q_{\mathrm{gs},\beta4}| < 0.5$. The cutoffs on the secondary structure CVs are to exclude states with significant C-terminal secondary structure, but our tests show that removing them does not significantly alter the calculations.

We would like to study dynamics of transitions between two disjoint states $A \subset O$ and $B \subset O$ that stay in $O$ the whole time (i.e., they avoid $\omega$). In our case, $A$ represents the gsKaiB state, $B$ represents the fsKaiB state, and $\omega$ represents a globally unfolded state. By computing the ratio of total reactive flux from paths that avoid the unfolded state $\omega$, $k^{O}_{\mathrm{gs}\to\mathrm{fs}}$, to the total reactive flux disregarding the unfolded state $\omega$, $k_{\mathrm{gs}\to\mathrm{fs}}$, we can obtain an estimate of the fraction of reactive trajectories which do or do not require unfolding as $1 - \frac{k^{O}_{\mathrm{gs}\to\mathrm{fs}}}{k_{\mathrm{gs}\to\mathrm{fs}}}$ or $\frac{k^{O}_{\mathrm{gs}\to\mathrm{fs}}}{k_{\mathrm{gs}\to\mathrm{fs}}}$, respectively.

To compute the reactive flux, we require the probability to first reach fsKaiB over gsKaiB, $q$, as well as the probability of having last come from gsKaiB over fsKaiB, $\bar{q}$ (16, 23, 24). We can estimate the $q$ and $\bar{q}$ for the paths that avoid the unfolded state using a modified equation

$$\mathcal{S}^{\tau}_{(\mathrm{A}\cup\mathrm{B}\cup\omega)^{\mathrm{c}}} q(x) = q(x) \text{ if } x \in O \backslash (A \cup B)$$

$$q(x) = 0 \text{ if } x \in A \cup \omega$$

$$q(x) = 1 \text{ if } x \in B$$

where $\mathcal{S}_S^{\tau} f(x) = \mathrm{E}\left[ f\left(X_{\min\{\tau, T^+(0)\}}\right) \mid X_0 = x \right]$ is the "stopped" transition operator and $T^+(0) = \min\{ s > 0 \mid X_s \in S^c\}$ is the first exit time from a set $S$ (22). One can define an equation similarly for $\bar{q}$ (see ref. (23) for further technical details). We then compute the total reactive flux following ref. (16) with a lag time of 200 frames (40,000 time steps) and with either $q$ or $1 - \bar{q}$ as the choice of reaction coordinate.

**Hydrogen-deuterium exchange simulations.**

HDX simulations were performed on the $\text{KaiB}^{\text{HDX}}$ mutant starting from the fsKaiB structure. Prolines were fixed in their initial isomerization state (i.e., *cis* for P63, P70, and P72) during the entire simulation. We ran T-REMD simulations with 48 replicas spaced between T = 0.78 and T = 1.02, each of length 2 million time steps and with attempts at Monte Carlo swaps every 10 frames. Following ref. (13) we compute the free energy of exchange for each residue *i* as

$$\Delta G_{\text{HDX,i}} = RT \ln\left(\frac{\langle w \cdot \text{PS}_i \rangle}{1 - \langle w \cdot \text{PS}_i \rangle}\right)$$

where $\text{PS}_i$ is a "protection state" value that roughly represents the competency of a given backbone hydrogen for exchange (between 0 and 1), and *w* is the weight assigned by MBAR reweighting. Angle brackets denote an ensemble average over all simulation frames and replicas.

Because Upside simulations of HDX effectively measure local stability, they correspond to the EX2 regime. This requires that $k_{\text{cl}} \gg k_{\text{chem}}$ where $k_{\text{cl}}$ and $k_{\text{chem}}$ are the closing and intrinsic exchange rates, respectively.

**All-atom simulations.**

Compared to the Upside simulations, all-atom MD simulations are computationally much more costly, as each time step represents only 2 fs of physical time, compared to 30 ps in Upside (a speed difference of 15,000x), even before accounting for the reduced number of degrees of freedom in the latter. On the other hand, the all-atom simulations provide a route to estimating quantitative free-energy differences. We simulated the temperature dependence of the free-energy difference between gsKaiB and fsKaiB as:

$$\Delta G_{FS-GS}(T) = \Delta G_{FS-GS}(T_0) + \Delta\Delta G_{FS-GS}(T - T_0)$$

$$\Delta\Delta G_{FS-GS}(T - T_0) = \Delta G_{FS}(T - T_0) - \Delta G_{GS}(T - T_0)$$

where $\Delta G_{FS-GS}(T)$ denotes the free energy of fold switching as a function of temperature, $T_0$ is an arbitrary reference temperature, and $\Delta\Delta G_{FS-GS}(T - T_0)$ represents the temperature-dependent shift in the free energy of fold switching. We focused on computing only $\Delta\Delta G_{FS-GS}(T - T_0)$, which is computationally much less costly because one only needs to simulate the free energy dependence on temperature for individual structures. To compute one such term, $\Delta G_{GS}(T - T_0)$, for example, one requires the Boltzmann factor that gives the relative probability of a structure at two temperatures, *i.e.*, $\frac{P(T_i)}{P(T_j)} = e^{-E\left(\frac{1}{k_b T_i} - \frac{1}{k_b T_j}\right)}$, which can be easily computed from the potential energies along the simulation trajectories. The Boltzmann factors can be used to compute the relative free energies of all temperature points using the MBAR method.

The structures of gsKaiB and fsKaiB used in the Upside simulations were also used for the all-atom simulations. The protonation states of both gsKaiB and fsKaiB were determined by H++ (25) at pH 7. The behavior of the protein systems was described by the AMBER-FB15 force field and the solvent was described by the TIP3P-FB water model. Previous studies have shown that this combination of force field and water model yields accurate melting curves for an α-helical peptide and β hairpin (26). After solvating the protein with 7210 waters in a 70 x 70 x 60 $Å^3$ box, eight $Cl^-$ ions were added to balance the charge of the system. The preparation of the MD simulations was done by *tleap* in Amber16 (27). Because the temperature dependence of free energy is proportional to the system size, we ensured that the protein sequence, and number of solvent molecules and ions were the same in both the gsKaiB and fsKaiB simulations.

A stepwise minimization process was performed to relax each system. First, the energy was minimized while subjecting the protein atoms to a harmonic energy restraint to their original positions. The side chains of the protein were optimized next, followed by an energy minimization without restraints. After the energy minimization process, a series of NVT simulations were carried using Langevin dynamics to heat the system to the targeted temperatures over 200 ps, followed by a NPT simulation in which the density was equilibrated at 1.0 atm using the Berendsen barostat over 200 ps. Finally, production MD simulations were carried out in the NPT ensemble. The simulations used the particle mesh Ewald (PME) treatment of electrostatic interactions with a real-space cutoff of 10 Å, as well as a cutoff of 10 Å for van der Waals interactions. A time step of 2 fs was used in the simulations and the SHAKE algorithm (28) was used to constrain the bonds to hydrogen atoms.

**Modeling the kinetics of fsKaiB$^{D91R}$-CI complex formation.**

We applied the following boundary conditions:

$$[gsKaiB^{D91R}] + [fsKaiB^{D91R}] + [fsKaiB^{D91R}\text{-}CI] = 1\ \mu M$$

$$[CI] + [fsKaiB^{D91R}\text{-}CI] = 10\ \mu M$$

The molar excess of CI ensures that the kinetics of formation of the fsKaiB$^{D91R}$-CI complex depends primarily on the properties of KaiB$^{D91R}$. We set $k_{on}$ to $10^4$ $h^{-1}$ $\mu M^{-1}$ so that fsKaiB$^{D91R}$-CI complexation is much faster than other rates. Regarding $k_{off}$, we previously determined it to be 0.03 $h^{-1}$ at 30 ºC for the fsKaiB$^{D91R}$-CI complex (29) and use that value here under the simplifying assumption that it is independent of temperature.

#### SUPPLEMENTARY FIGURES

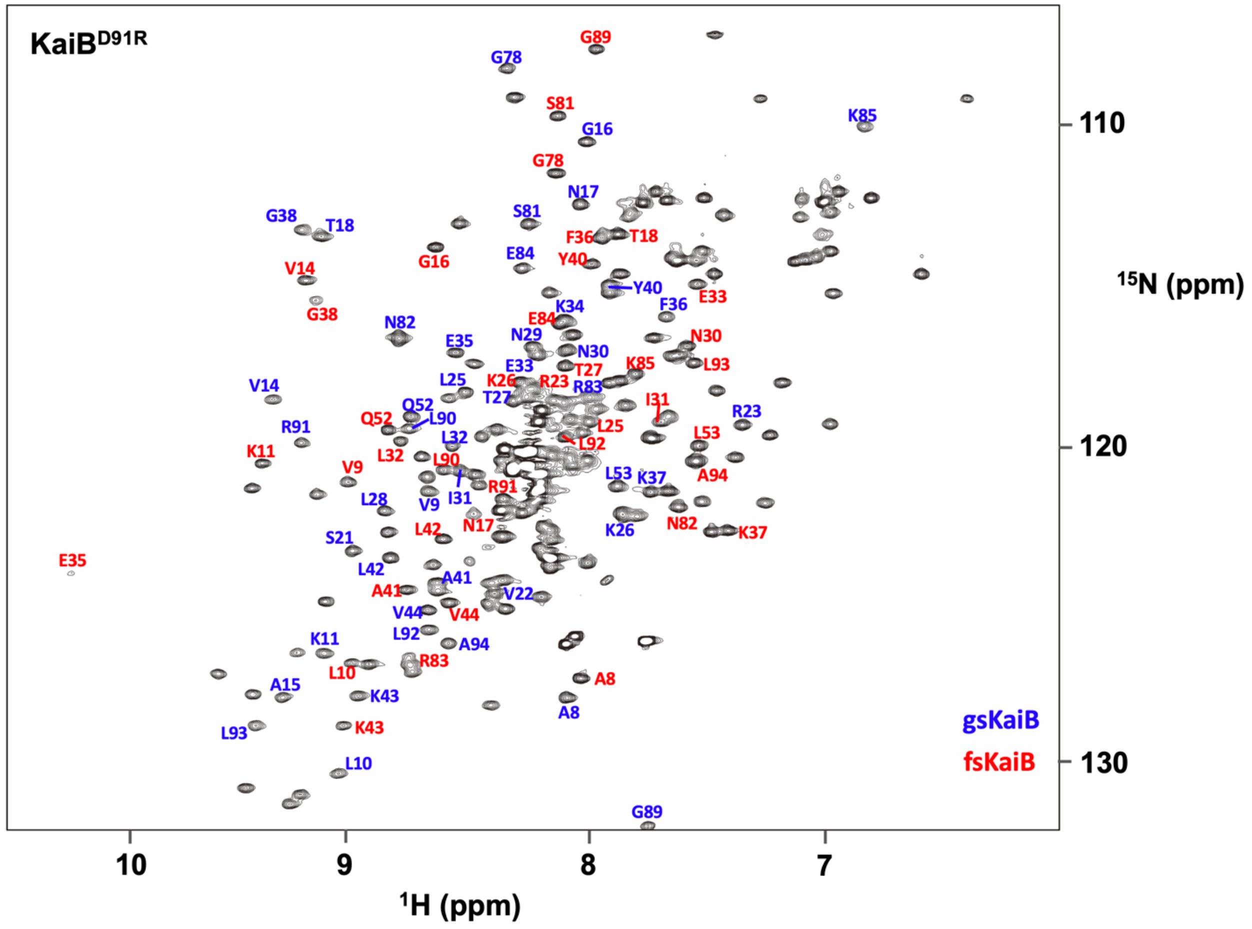


**Figure S1. Assigned peaks in the $^{1}H$-$^{15}N$ HSQC spectrum of KaiB$^{D91R}$.** The spectrum was acquired at a sample temperature of 25 ºC. Assigned peaks for gsKaiB and fsKaiB are indicated with blue and red labels, respectively. Chemical-shift assignments were determined by Chang et al., 2015 (29).

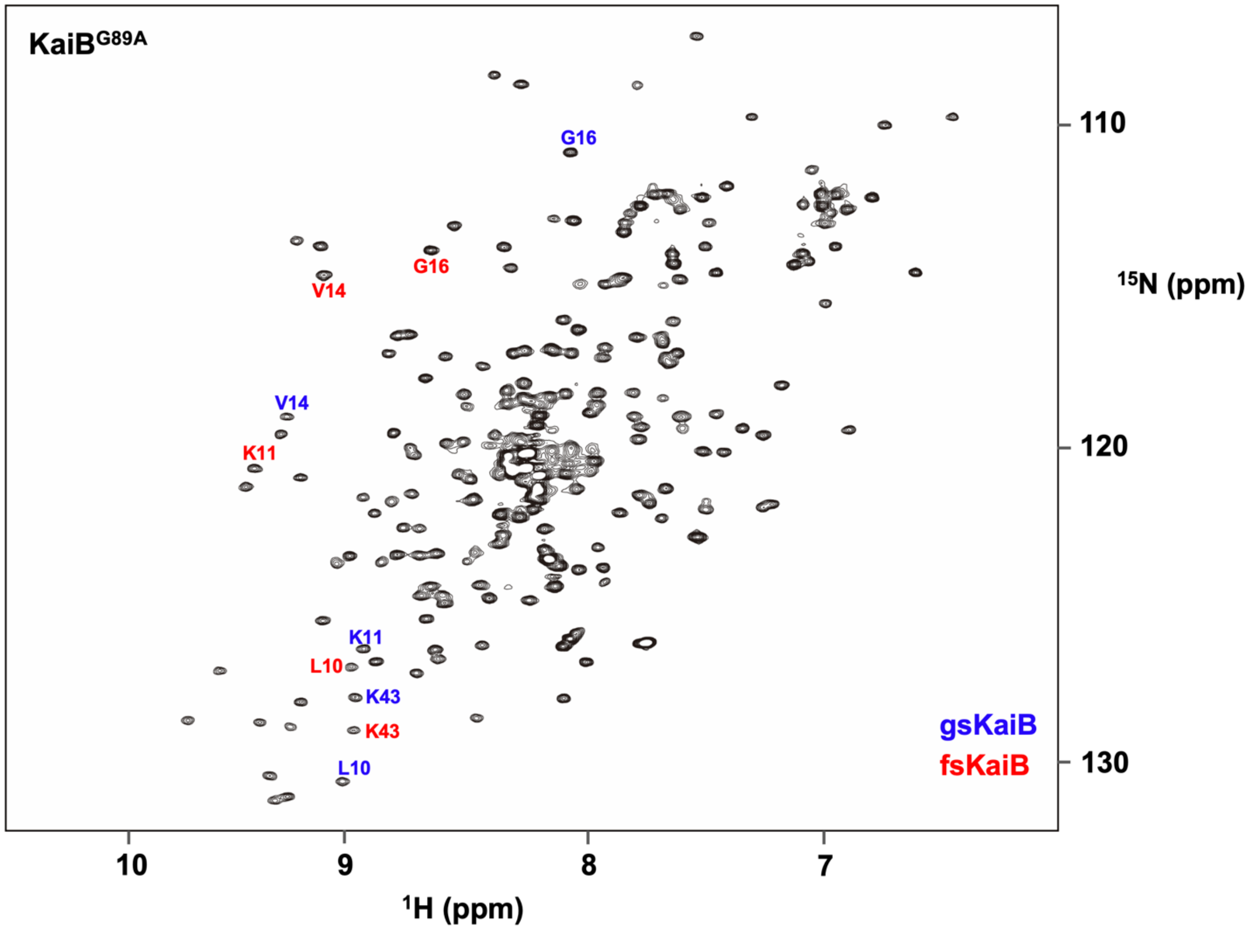


**Figure S2. Assigned peaks in the $^{1}$H-$^{15}$N HSQC spectrum of KaiB$^{G89A}$.** The spectrum was acquired at a sample temperature of 25 ºC. Assigned peaks for gsKaiB and fsKaiB are indicated with blue and red labels, respectively. Chemical-shift assignments were determined by Chang et al., 2015 (29).

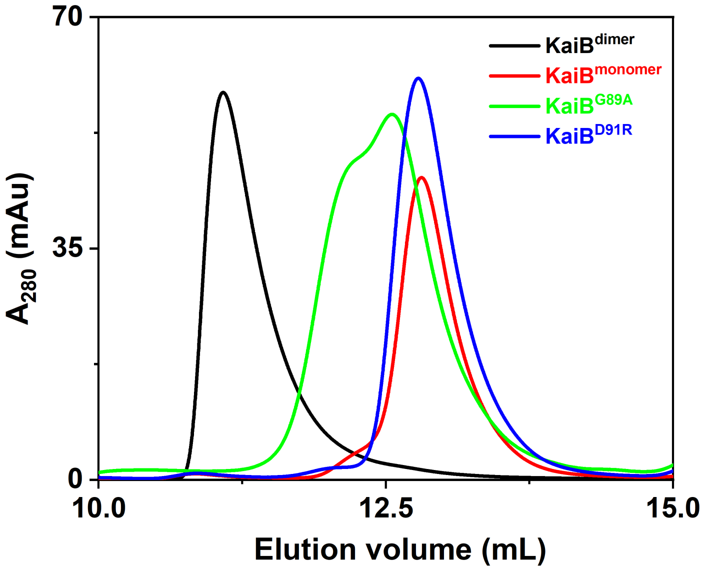


**Figure S3. Size-exclusion chromatograms of KaiB constructs at room temperature.** KaiB$^{dimer}$ (KaiB 1-94 Y8A Y94A (black)), KaiB$^{monomer}$ (KaiB 1-94 Y8A G89A D91R Y94A (red)), KaiB$^{G89A}$ (KaiB 1-94 Y8A G89A Y94A (green)), and KaiB$^{D91R}$ (KaiB 1-94 Y8A, D91R, Y94A (blue)). KaiB$^{monomer}$ and KaiB$^{dimer}$ were previously determined to be monomeric and dimeric, respectively (29).

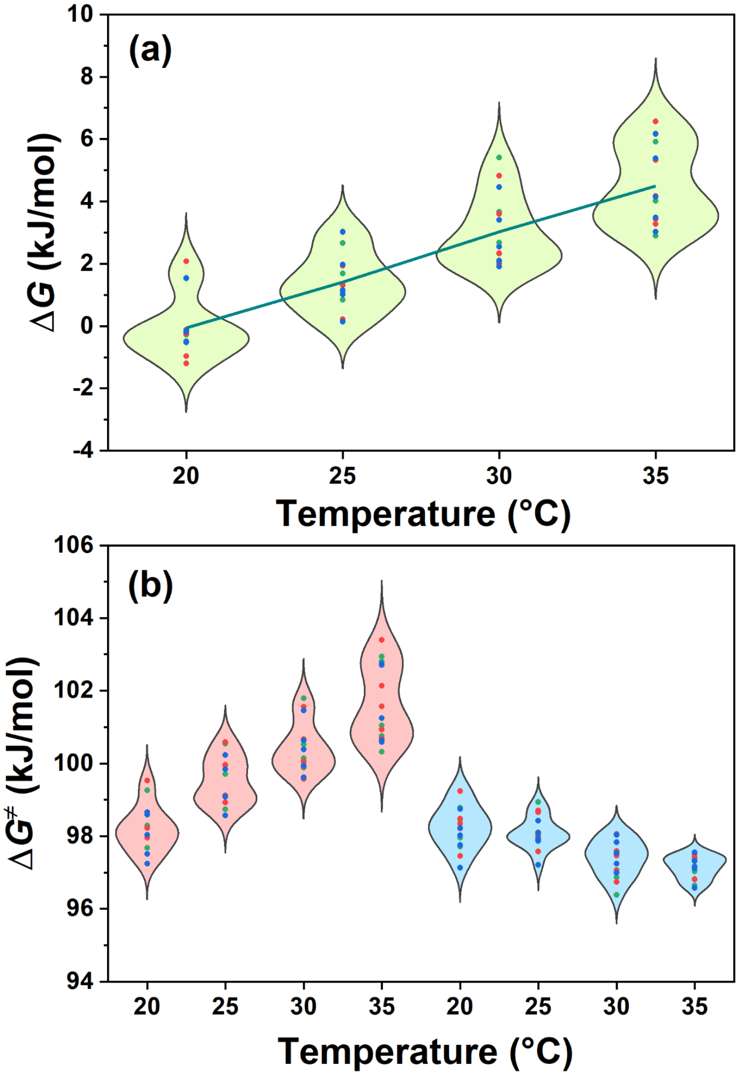


**Figure S4. Temperature dependence of the gsKaiB$^{G89A}$ ⇌ fsKaiB$^{G89A}$ reaction. (a)** Residue-specific Δ*G* values for gsKaiB ⇌ fsKaiB are plotted as a function of temperature. Red, green, and blue points are used to distinguish data from three replicates and are superimposed on distribution diagrams for each temperature sampled. The blue line is a fit to mean values as a function of temperature. **(b)** $\Delta G^{‡}_{gs\rightarrow fs}$ (pink) and $\Delta G^{‡}_{gs\leftarrow fs}$ (blue) values as a function of temperature. Red, green, and blue data points represent different replicate data sets.

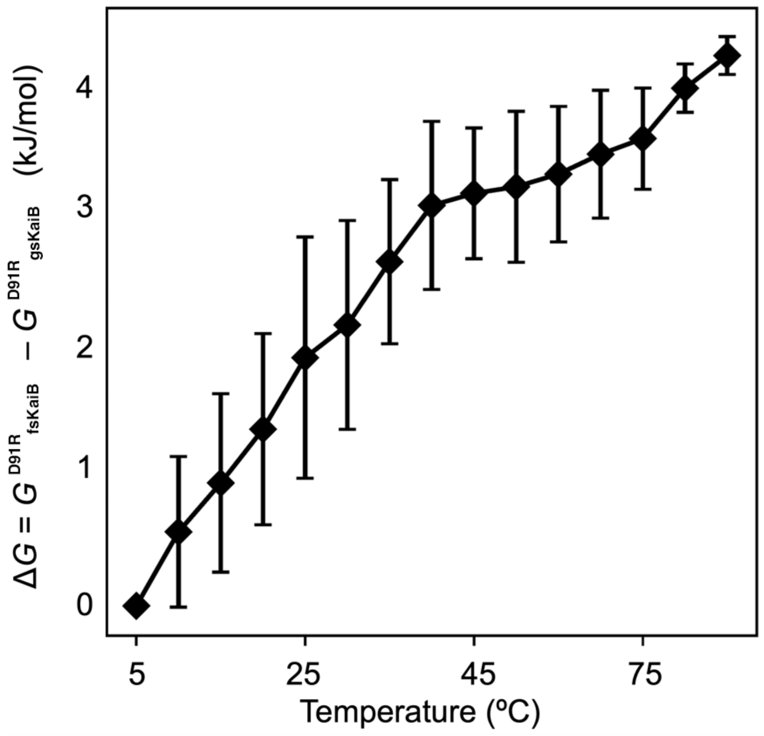


**Figure S5. Temperature dependence of the free-energy difference, Δ$G$ = $G^{D91R}{}_{fsKaiB}$ – $G^{D91R}{}_{gsKaiB}$, from all-atom molecular dynamics simulations.** Error bars were computed from repeating the analysis twice, once using only the first half of the trajectory and a second time using only the second half.

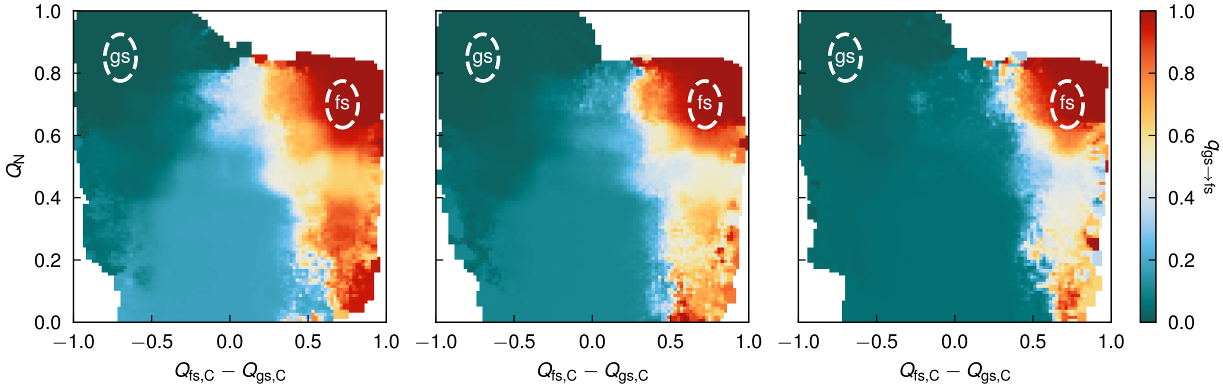


**Figure S6. Fold switching progress for an Upside model of KaiB$^{D91R}$.** We show the probability of reaching fsKaiB before gsKaiB as a function of the fraction of N-terminal contacts ($Q_N$) and the difference in C-terminal contacts to the gsKaiB and fsKaiB stuctures ($Q_{fs,C} - Q_{gs,C}$). From left to right, the Upside temperature is increasing.

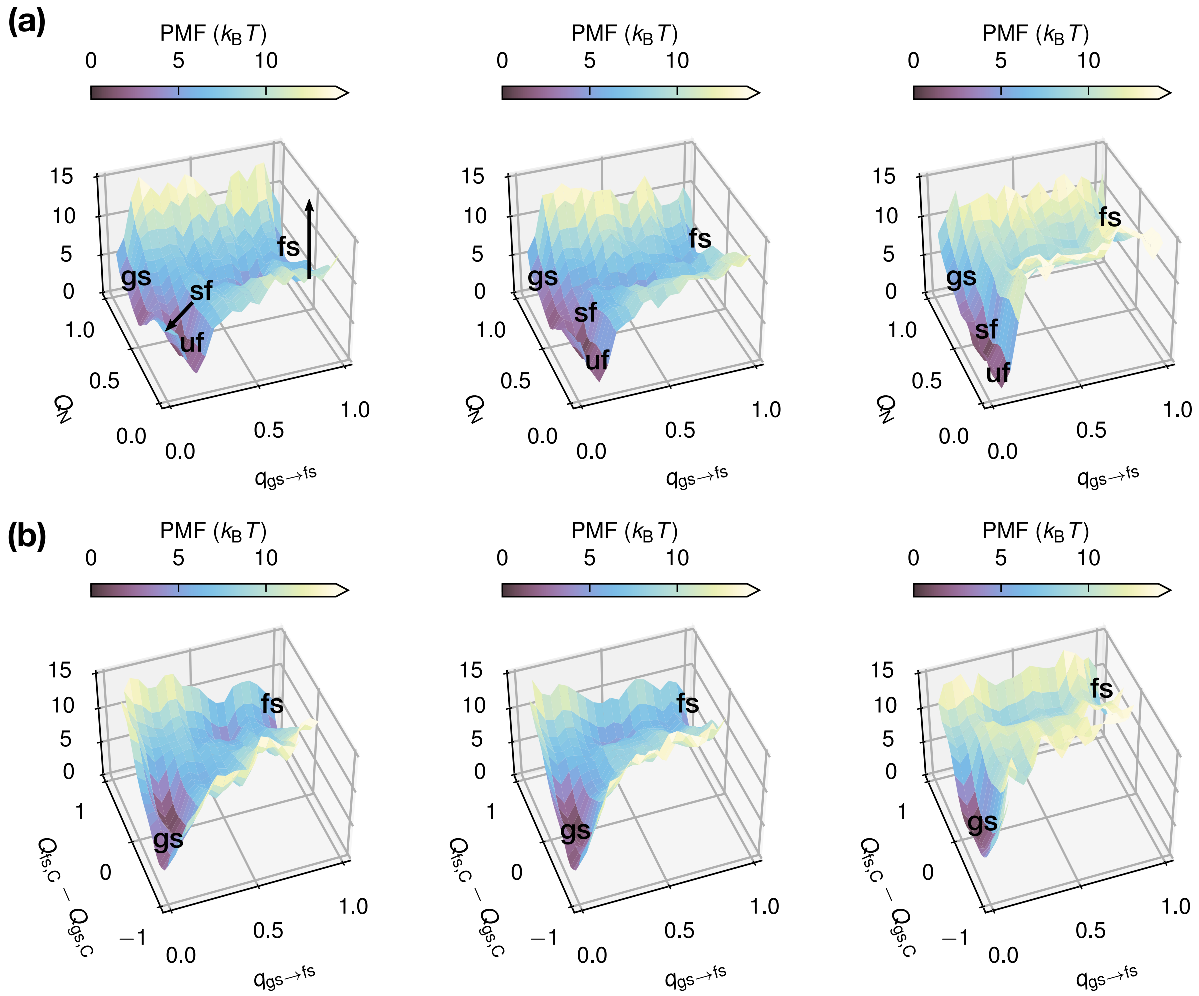


**Figure S7. Free energy profiles of fold switching for an Upside model of KaiB$^{D91R}$.** Potentials of mean force as a function of the probability of proceeding to fsKaiB over gsKaiB, *q*, and **(a)** the fraction of native contacts in the N-terminal half of the protein ($Q_N$), and **(b)** the difference in the fractions of C-terminal contacts between the gsKaiB and fsKaiB states ($Q_{fs,C} – Q_{gs,C}$). The "sf" (semi-folded) and "uf" (unfolded) states correspond approximately to the metastable intermediates at (0.0, 0.4) and (0.0, 0.0) in the plots in **Fig. 4**. The arrows depict the movement of the semi-folded and unfolded states towards gsKaiB and the increasing free energy of fsKaiB as temperature increases. From left to right, the Upside temperature is increasing.

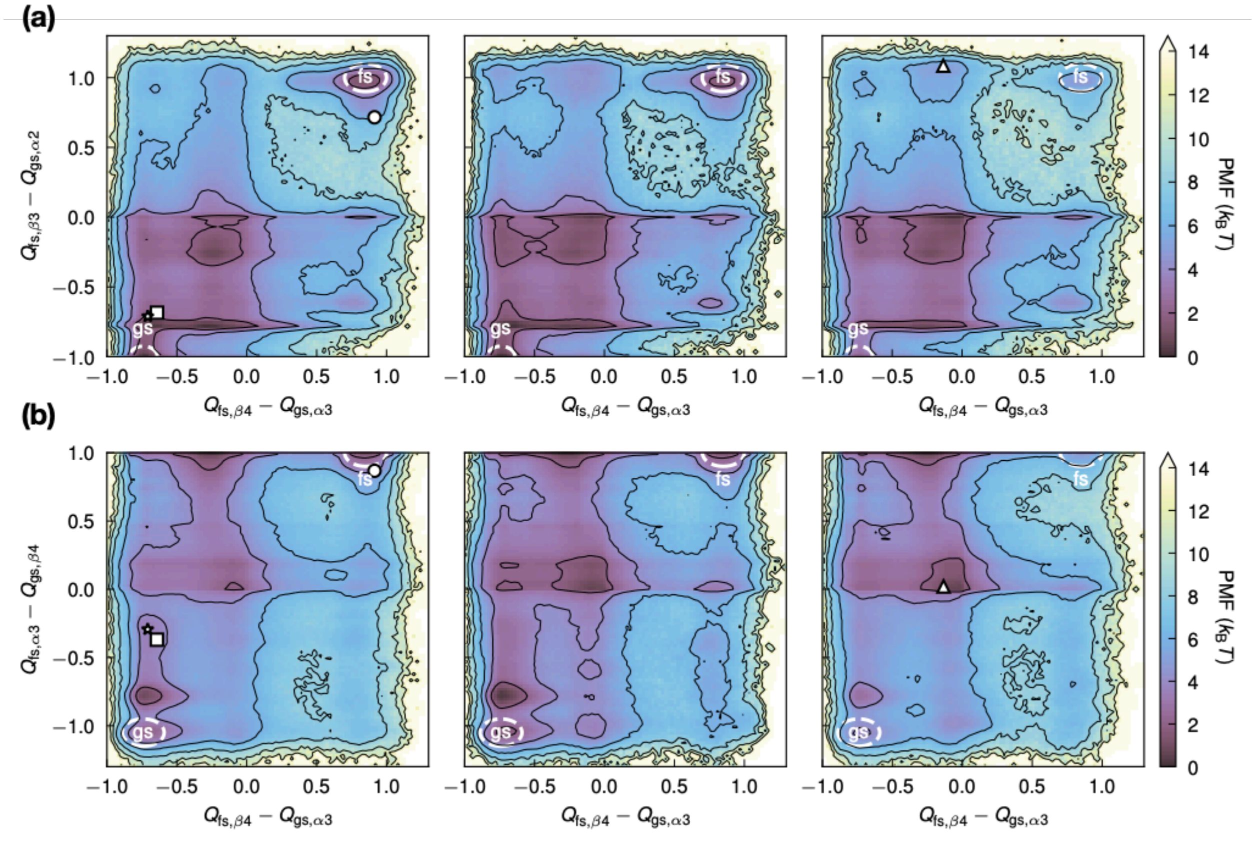


**Figure S8. Free-energy profiles of the fold-switching secondary structure for an Upside model of KaiB$^{D91R}$.** Potentials of mean force as a function of the **(a)** $Q_{fs,\beta4} - Q_{gs,\alpha3}$ and $Q_{fs,\alpha3} - Q_{gs,\beta4}$ CVs and **(b)** $Q_{fs,\beta4} - Q_{gs,\alpha3}$ and $Q_{fs,\beta3} - Q_{gs,\alpha2}$ CVs. Approximate location of the intermediate structures shown in **Fig. 4a** are marked with the corresponding symbols. From left to right, the Upside temperature is increasing. Contour lines are drawn every 2 $k_B T$.

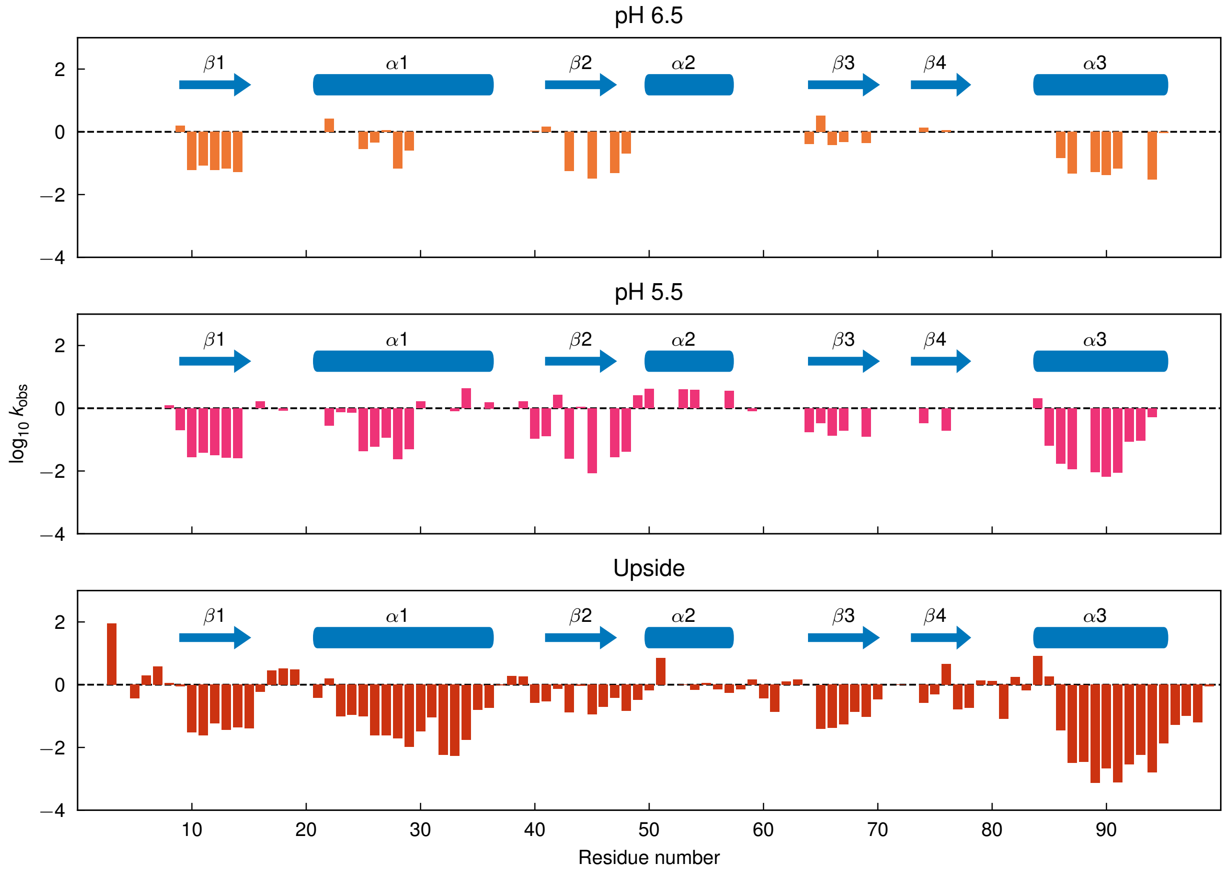


**Figure S9. Observed and computed hydrogen-deuterium exchange rates of KaiB$^{HDX}$.** From top to bottom, $\log_{10}$ of $k_{obs}$ rates (in $h^{-1}$) for experiments at pH 6.5 and pH 5.5, and from Upside coarse-grained simulations. Upside simulations correspond to T = 0.91 and are computed by converting stabilities (free energies) using the experimentally measured $k_{chem}$ values (see supplementary discussion) at pH 5.5 with the relation $k_{obs} = k_{chem}/PF$, where $PF = 1 + \exp(\Delta G_{HDX}/RT)$, assuming a temperature conversion of 0.85 ↔ 298 K.

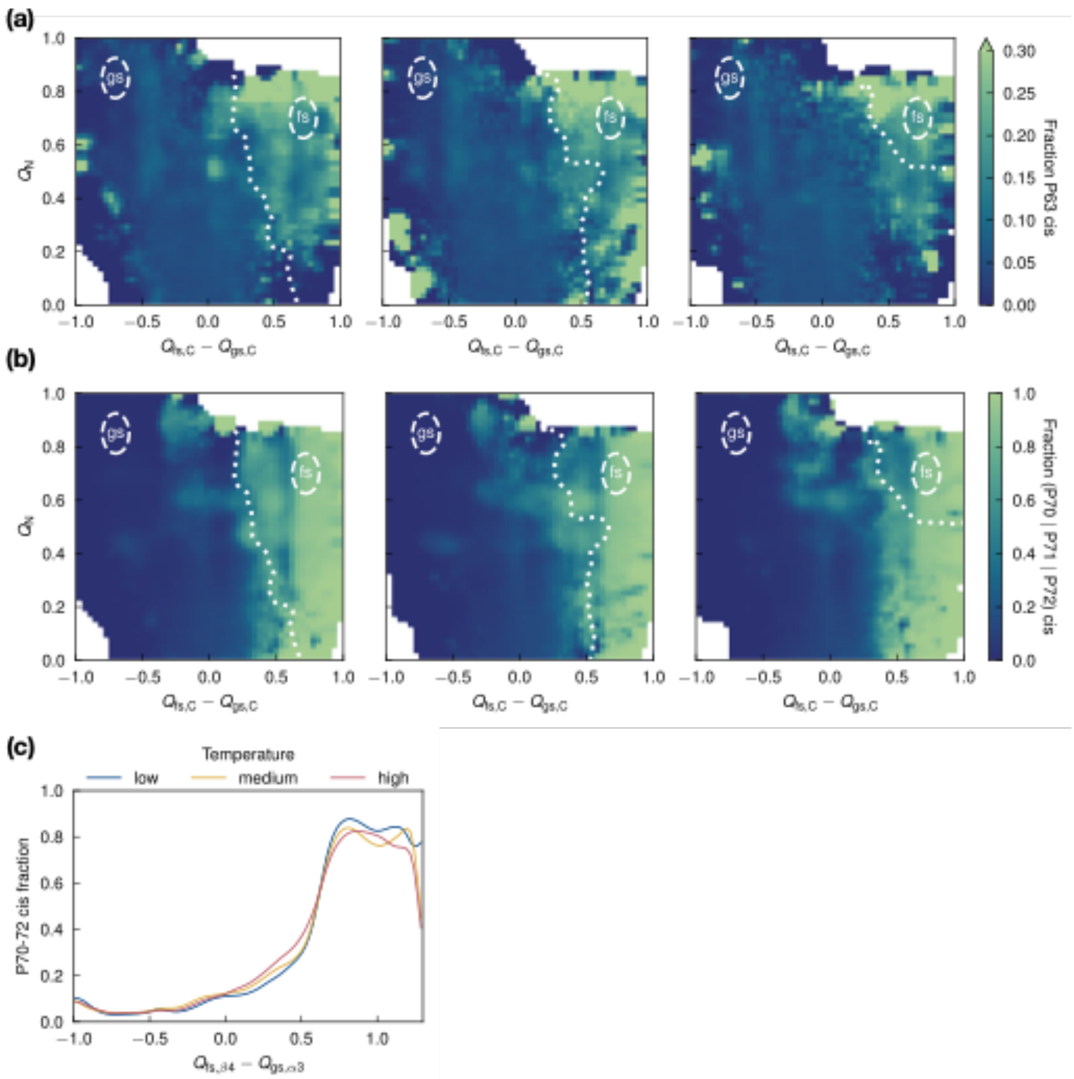


**Figure S10. Role of isomerization of prolines in Upside simulations of KaiB$^{D91R}$ fold switching.** Average fraction of **(a)** P63 and **(b)** any of P70–P72 in *cis* as a function of the fraction of N-terminal contacts ($Q_N$) and the difference in C-terminal contacts to the fs and gs structures ($Q_{fs,C} - Q_{gs,C}$). From left to right, the Upside temperature is increasing. The approximate location of the $q$ = 0.5 surface is marked as a dashed line. **(c)** Average fraction of any of P70–P72 in *cis* as a function of $Q_{fs,\beta4} - Q_{gs,\alpha3}$.

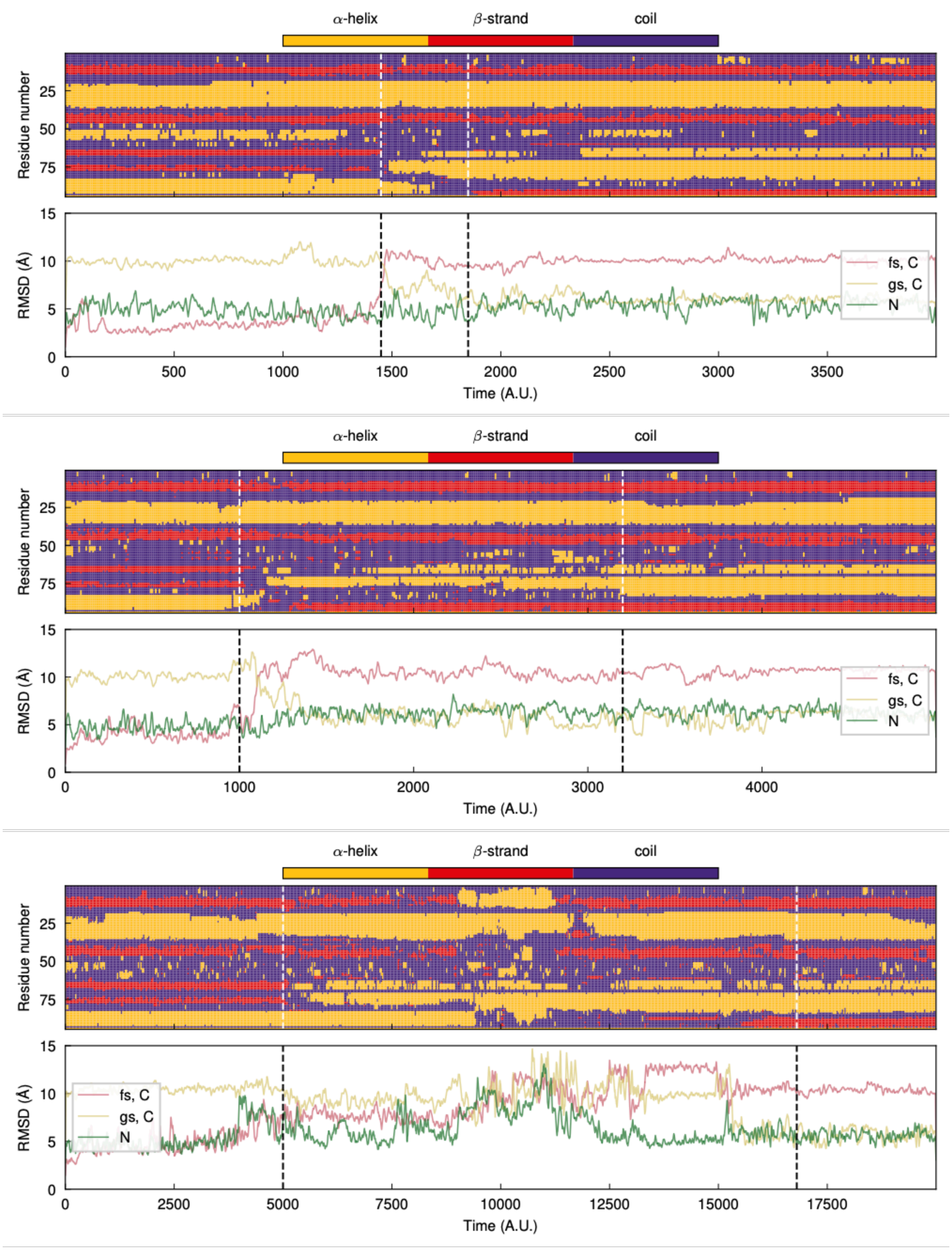


**Figure S11. Proline isomerization is sufficient to induce fold-switching in an**

**Upside model of KaiB$^{D91R}$**. Upside simulations starting from fsKaiB with P63, P70, and P72 fixed in *trans*. (top) Secondary structure assigned via DSSP (30) and (bottom) RMSD to either the C-terminal half of fsKaiB, the C-terminal half of gsKaiB, or the N-terminal half (**Table S1**) during three other successful fold-switching simulations from fsKaiB to gsKaiB. RMSDs are plotted as a moving average over 20 frames. The dotted lines indicate the approximate start and end times of the fold-switching event.

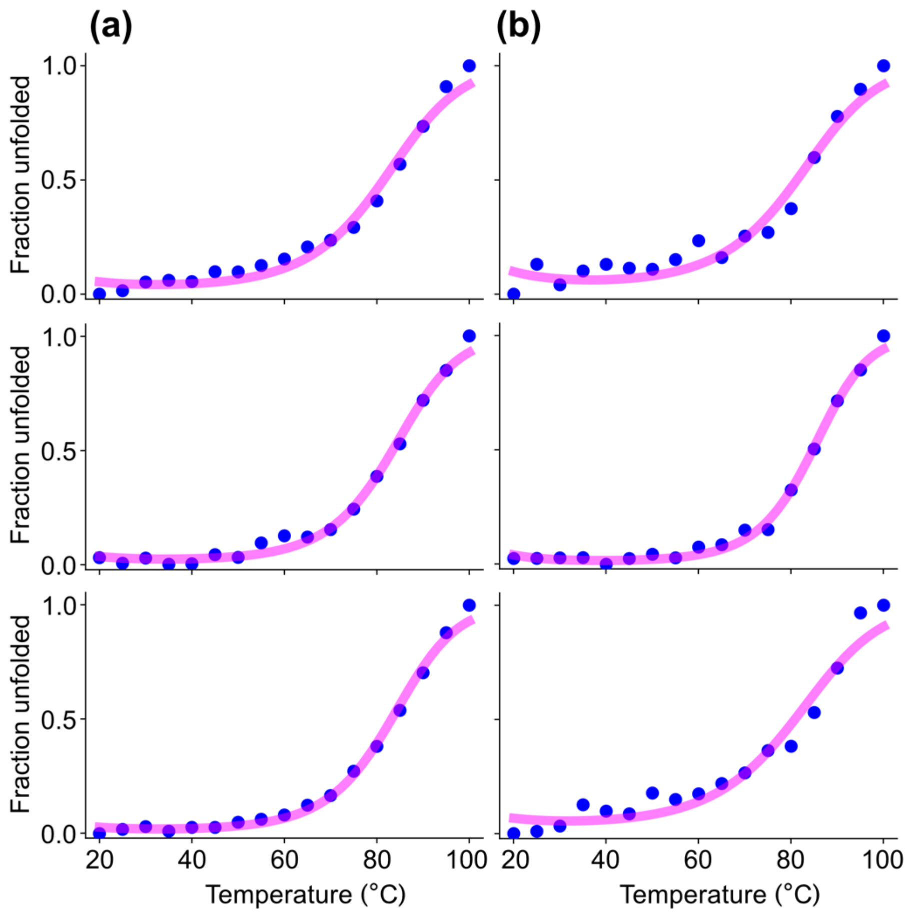


**Figure S12. Thermal denaturation profiles for KaiB$^{HDX}$ at (a) pH 5.5 and (b) pH 6.5.** Three replicate experiments were carried out for both pH values on a Jasco J-1500 CD spectrometer. Fits (pink curves) to the data (blue points) yielded melting points of 83 ± 1 ºC and 82 ± 1 ºC for pH 5.5 and pH 6.5, respectively. The accuracy of these melting points is approximate as full denaturation of KaiB$^{HDX}$ was not achieved over this range of temperatures. The concentrations of KaiB$^{HDX}$ were 14 μM and 12 μM at pH 5.5 and pH 6.5, respectively, in 20 mM sodium phosphate and 100 mM NaCl, titrated with HCl to adjust pH values. Measurements were done at a wavelength of 220 nm at 5 ºC intervals from 20 ºC to 100 ºC (17 data points total) with a temperature ramp of 1 ºC /min. Each data point represents the average value of seven readings at 220 nm.

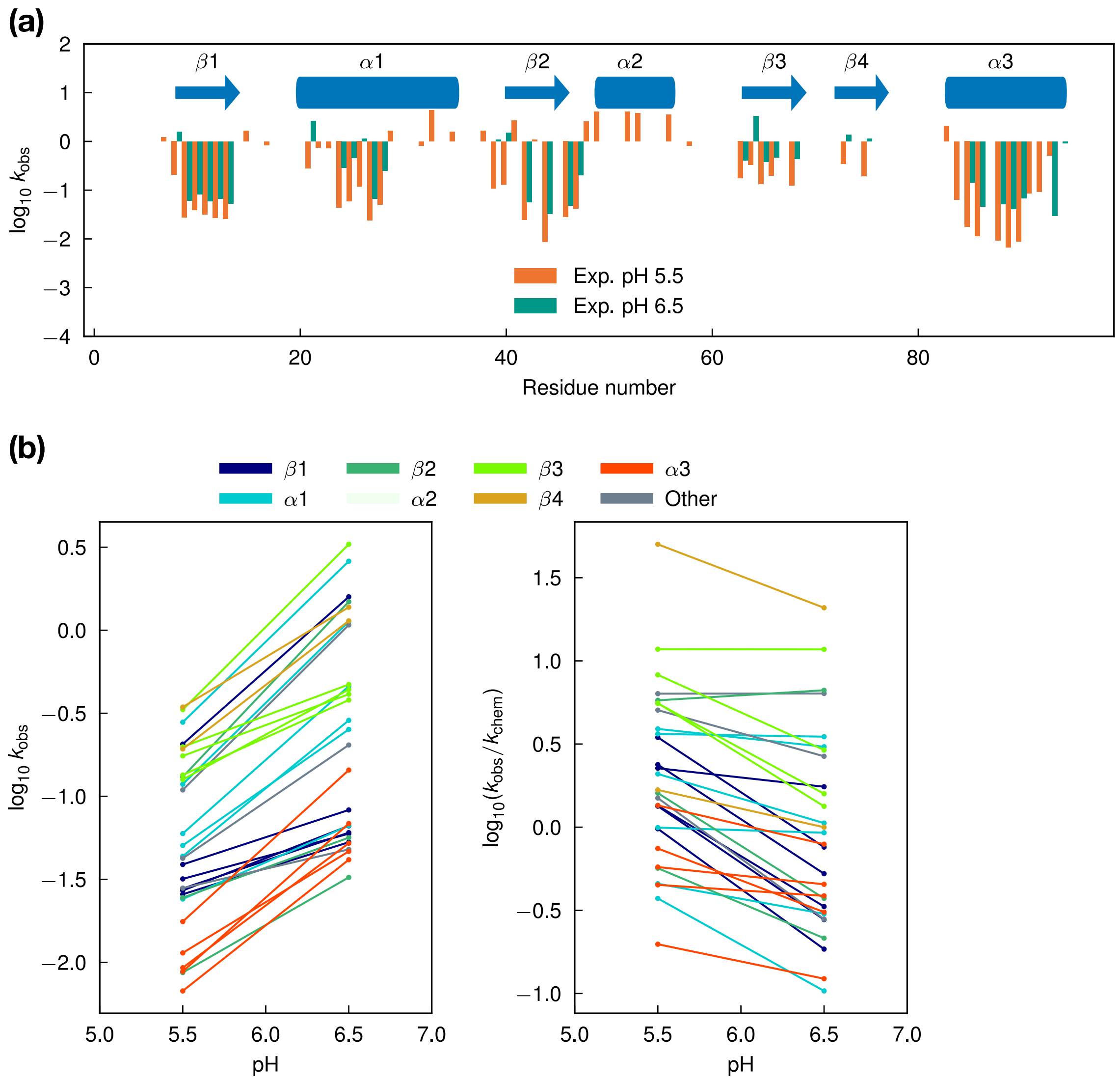


**Figure S13. HDX rates for KaiB$^{HDX}$ as a function of pH. (a)** $\log_{10}$ of $k_{obs}$ (in $h^{-1}$) for residues measured at pH 5.5 and 6.5. **(b)** Left, $\log_{10}k_{obs}$, and right, $\log_{10}(k_{obs}/k_{chem})$ at same pH values and colored by fsKaiB secondary structure. In the plot of $k_{obs}$ and the plot of $k_{obs}/k_{chem}$, a slope of 1 and 0 respectively, can be attributable to a 10-fold increase in $k_{obs}$ with a one unit increase in pH matching a similar 10-fold increase in $k_{chem}$, and are the expected slopes in the EX2 limit.

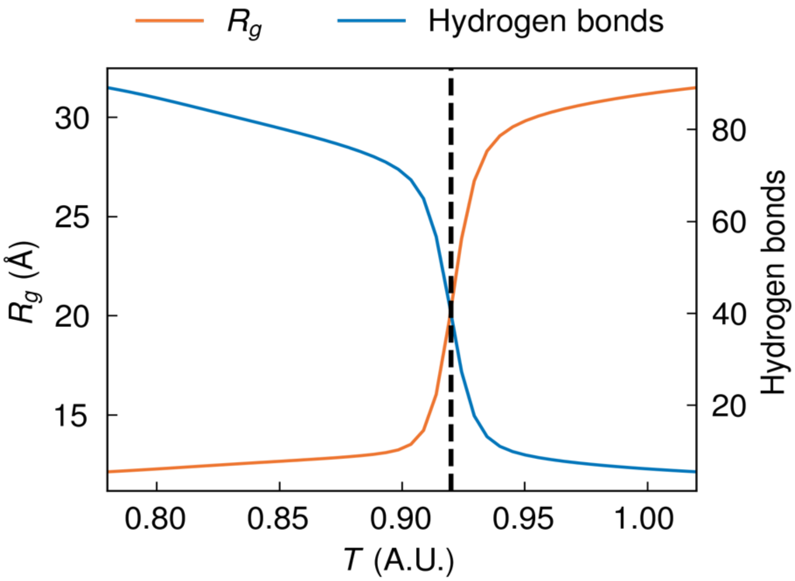


**Figure S14. Melting curve for Upside model of KaiB$^{D91R}$.** The black dashed line indicates the estimated melting temperature of T = 0.92. $R_g$ is the radius of gyration.

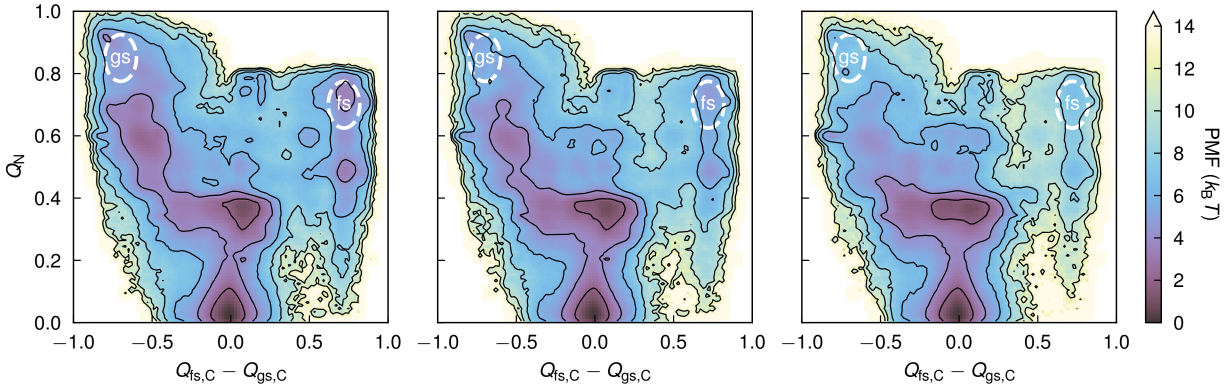


**Figure S15. Free-energy profile of KaiB$^{D91R}$ from T-REMD Upside simulations.** Potential of mean force as a function of the fraction of N-terminal contacts ($Q_N$) and the difference in the fractions of C-terminal contacts in the fs and gs structures ($Q_{fs,C} – Q_{gs,C}$). From left to right, the temperature is increasing. Contour lines are drawn every 2 $k_B T$.

### SUPPLEMENTARY TABLES

**Table S1. Residue definitions, in terms of amino acid numbers, for the secondary structure elements and N/C-termini in KaiB$^{D91R}$.**

| Name | gs | fs |
|---|---|---|
| N-terminal half | 1–50 | 1–50 |
| C-terminal half | 51–94 | 51–94 |
| $\beta 3_{gs} \rightleftarrows \alpha 2_{fs}$ | 59–61 | 50–57 |
| $\alpha 2_{gs} \rightleftarrows \beta 3_{fs}$ | 62–67 | 64–69 |
| $\alpha 3_{gs} \rightleftarrows \beta 4_{fs}$ | 72–81 | 73–77 |
| $\beta 4_{gs} \rightleftarrows \alpha 3_{fs}$ | 88–94 | 84–94 |

**Table S2. Equilibrium population fraction of the *cis* isomerization state for selected prolines in Upside simulations of KaiB$^{D91R}$**

| % cis | P63 | P70 | P71 | P72 |
|---|---|---|---|---|
| **gsKaiB*** | 0.0 | 0.1 | 0.0 | 5.6 |
| **fsKaiB*** | 28.7 | 8.2 | 89.9 | 3.8 |

*The gsKaiB and fsKaiB states were defined as in **Table S4**.

**Table S3. List of KaiB variants.**

<table>
<tr><td colspan="2">KaiB$^{G89A}$ for NMR temperature-jump experiments and SEC</td></tr>
<tr><td>Variant</td><td>FLAG-TeKaiB-1-94-Y8A-Y94A-G89A-FLAG</td></tr>
<tr><td>Construct</td><td>pET-28b-SUMO-FLAG-TeKaiB-1-94-Y8A-Y94A-G89A-FLAG</td></tr>
<tr><td>Protein sequence</td><td>DYKDDDDKMAPLRKTAVLKLYVAGNTPNSVRALKTLNNILEKEFKGVYA<br>LKVIDVLKNPQLAEEDKILATPTLAKVLPPPVRRIIGDLSNREKVLIALDLLA<br>DYKDDDDK</td></tr>
<tr><td colspan="2">KaiB$^{D91R}$ for NMR temperature-jump experiments and SEC</td></tr>
<tr><td>Variant</td><td>FLAG-TeKaiB-1-94-Y8A-Y94A-D91R-FLAG</td></tr>
<tr><td>Construct</td><td>pET-28b-SUMO-FLAG-TeKaiB-1-94-Y8A-Y94A-D91R-FLAG</td></tr>
<tr><td>Protein sequence</td><td>DYKDDDDKMAPLRKTAVLKLYVAGNTPNSVRALKTLNNILEKEFKGVYA<br>LKVIDVLKNPQLAEEDKILATPTLAKVLPPPVRRIIGDLSNREKVLIGLRLLA<br>DYKDDDDK</td></tr>
<tr><td colspan="2">KaiB$^{Dimer}$ for SEC</td></tr>
</table>

| | |
|---|---|
| Variant | FLAG-*Te*KaiB-1-94-Y8A-Y94A -FLAG |
| Construct | pET-28b-SUMO-FLAG-*Te*KaiB-1-94-Y8A-Y94A -FLAG |
| Protein sequence | DYKDDDDKMAPLRKTAVLKLYVAGNTPNSVRALKTLNNILEKEFKGVYA LKVIDVLKNPQLAEEDKILATPTLAKVLPPPVRRIIGDLSNREKVLIGLDLLA DYKDDDDK |
| **KaiB$^{Monomer}$ for SEC** | |
| Variant | FLAG-*Te*KaiB-1-94-Y8A-Y94A-G89A-D91R-FLAG |
| Construct | pET-28b-SUMO-FLAG-*Te*KaiB-1-94-Y8A-Y94A-G89A-D91R-FLAG |
| Protein sequence | DYKDDDDKMAPLRKTAVLKLYVAGNTPNSVRALKTLNNILEKEFKGVYA LKVIDVLKNPQLAEEDKILATPTLAKVLPPPVRRIIGDLSNREKVLIALRLLA DYKDDDDK |
| **KaiB$^{HDX}$ for Hydrogen-deuterium exchange (HDX) experiments** | |
| Variant | *Te*KaiB-1-99-Y8A-Y94A-P71A-G89A-D91R |
| Construct | pET-28b-SUMO-*Te*KaiB-1-99-Y8A-Y94A-P71A-G89A-D91R |

| | |
|---|---|
| Protein sequence | MAPLRKTAVLKLYVAGNTPNSVRALKTLNNILEKEFKGVYALKVIDVLKN PQLAEEDKILATPTLAKVLPPPVRRIIGDLSNREKVLIGLRLLA |

*Te*: abbreviation for *Thermosynechococcus elongatus BP-1*

FLAG: a tag for improving solubility of KaiB. The sequence is DYKDDDDK

SUMO: a protein fusion tag for enhancing the expression and solubility of the target recombinant protein.

**Table S4. State definitions of gsKaiB and fsKaiB for DGA calculations.**

| CV | gs cutoff | fs cutoff |
|---|---|---|
| $Q_{fs,\alpha2} - Q_{gs,\beta3}$ | < –0.50 | – |
| $Q_{fs,\beta3} - Q_{gs,\alpha2}$ | < –0.75 | > 0.85 |
| $Q_{fs,\beta4} - Q_{gs,\alpha3}$ | < –0.55 | > 0.65 |
| $Q_{fs,\alpha3} - Q_{gs,\beta4}$ | < –0.90 | > 0.80 |
| $Q_N$ | > 0.70 | > 0.65 |
| $Q_s$ ($s$ = gs or fs) | > 0.62 | > 0.61 |
| $Q_{s,C}$ ($s$ = gs or fs) | > 0.65 | > 0.75 |
| RMSD to $s$ (nm) | < 0.45 | < 0.35 |
